# Minimum entropy framework identifies a novel class of genomic functional elements and reveals regulatory mechanisms at human disease loci

**DOI:** 10.1101/2023.06.11.544507

**Authors:** Michael J. Betti, Melinda C. Aldrich, Eric R. Gamazon

**Affiliations:** Vanderbilt University Medical Center, Nashville, TN; Clare Hall, University of Cambridge, Cambridge, England

## Abstract

We introduce CoRE-BED, a framework trained using 19 epigenomic features in 33 major cell and tissue types to predict cell-type-specific regulatory function. CoRE-BED identifies nine functional classes *de-novo*, capturing both known and new regulatory categories. Notably, we describe a previously undercharacterized class that we term Development Associated Elements (DAEs), which are highly enriched in cell types with elevated regenerative potential and distinguished by the dual presence of either H3K4me2 and H3K9ac (an epigenetic signature associated with kinetochore assembly) or H3K79me3 and H4K20me1 (a signature associated with transcriptional pause release). Unlike bivalent promoters, which represent a transitory state between active and silenced promoters, DAEs transition directly to or from a non-functional state during stem cell differentiation and are proximal to highly expressed genes. CoRE-BED’s interpretability facilitates causal inference and functional prioritization. Across 70 complex traits, distal insulators account for the largest mean proportion of SNP heritability (∼49%) captured by the GWAS. Collectively, our results demonstrate the value of exploring non-conventional ways of regulatory classification that enrich for trait heritability, to complement existing approaches for *cis*-regulatory prediction.

## Introduction

Since the advent of the genome-wide association study (GWAS) methodology^1^, there has been remarkable growth in the number of population-level genetic studies covering large swaths of the human phenome. However, the GWAS approach has been hindered by a consistent bottleneck: the functional ambiguity of most single nucleotide polymorphisms (SNPs) identified through the methodology. Ninety percent of GWAS-identified variants are predominantly located in non-coding regions of the genome and do not directly impact the coding sequence of proteins.^2^ Consequently, determining the underlying mechanisms for the observed associations remains a challenge. Despite the accelerated pace of genetic discoveries from GWAS, less than 1% of these associations have been functionally confirmed through follow-up studies.^3^ This discrepancy underscores the need for improved *in silico* approaches to help bridge the knowledge gap, as the scale of the problem makes experimental validation infeasible.

Integrating GWAS results with omics information can help to elucidate the function of non-coding variants^2,4^. In addition to the ENCODE Consortium’s comprehensive registry of candidate *cis*-regulatory elements (cCREs) across over 1,500 cell types^5^, prediction frameworks such as Activity-by-Contact^6^ and ChromHMM^7,8^ integrate rich functional genomics datasets to predict regulatory function in non-coding regions of the genome.

Here, we introduce CoRE-BED, a decision tree-based framework that has been trained on 19 epigenomic features to learn regulatory classes. In contrast to deterministic models of genome regulation, which assert that individual regulatory classes are characterized by a single specific combination of regulatory marks, CoRE-BED regulatory elements are defined through multiple paths; there is no *a priori* definition of feature combinations for a given class. Thus, the distribution of epigenomic marks within each class resembles a spectrum rather than a set of discrete signatures.

Because CoRE-BED utilizes a decision tree-based architecture, the model’s functional predictions provide high interpretability^9^, offering biological insights into genome structure and function. We demonstrate that the functional classes learned *de novo* by CoRE-BED provide both expected classifications (active promoters and insulators) and novel ways of grouping regulatory categories, including, for example, a class that we term Development Associated Elements (DAEs). Additionally, we show that CoRE-BED is an effective tool for GWAS variant prioritization, with the ability to functionally prioritize a set of SNPs enriched for trait heritability. In summary, the CoRE-BED framework is characterized by high interpretability and practical applicability, providing robust prediction of regulatory function and identifying high-priority SNPs for post-GWAS analyses.

## Results

### CoRE-BED framework and regulatory predictions

We present CoRE-BED, a decision tree framework for predicting cell and tissue type-specific regulatory elements across the genome. Despite the availability of more sophisticated deep learning-based approaches, tree-based models still hold their own against neural networks when working with tabular data^10^. An additional motivation for choosing a decision tree was its high interpretability. The decision tree structure was learned from training data, generating a classifier (i.e., membership in a class of regulatory elements) whose generalizability was evaluated in test data (Figure 1). Each internal node represents an observation that captures a physical property of chromatin, nucleosome positioning, or DNA accessibility; each leaf or terminal node is a class of regulatory elements; and each link is a decision. To determine the optimal tree structure, we considered the information gain, calculated using Shannon entropy^11^, at each node split. Classification was performed by traversing down the tree hierarchy from the root node to the terminal node. The benefits of the decision tree framework include improved interpretability, ease of implementation (e.g., scaling and normalization not a prerequisite), and a non-parametric methodology (requiring no distributional assumptions).

**Figure 1.**
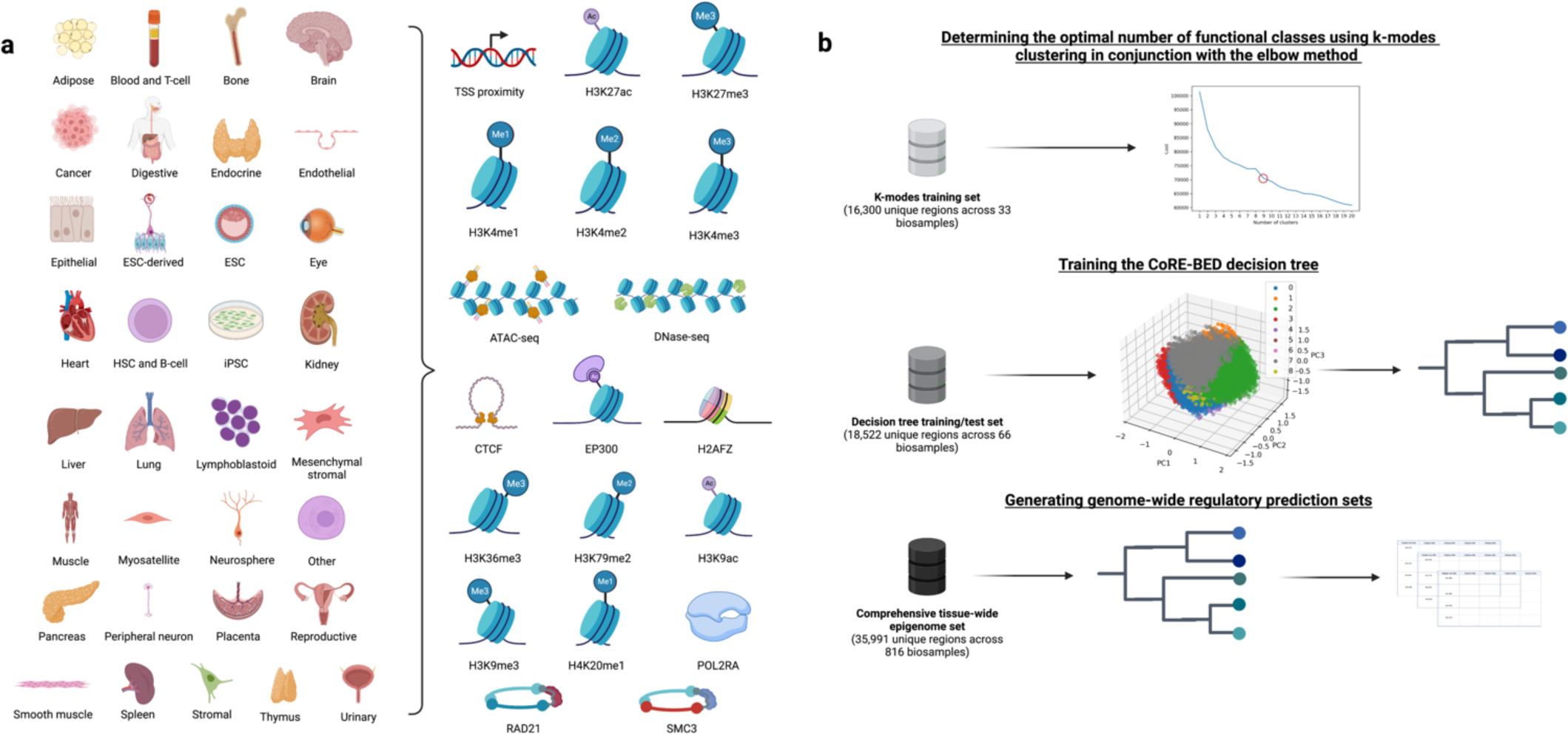
The CoRE-BED framework. **a,** The CoRE-BED training set was composed of 19 distinct epigenomic features representing 33 major categories of cell and tissue types. **b,** Development of the framework consisted of three steps. First, we determined the optimal number of clusters making up the data using k-modes clustering in conjunction with the elbow method. Next, using the learned k = 9, k-modes clustering was performed on an independent dataset, i.e., the training set, for the decision tree implementation. Finally, the trained decision tree was used to predict genome-wide functional classes for 816 individual biosamples encompassing the 33 major cell and tissue categories.

The decision tree model was trained using 19 epigenomic features from 33 major cell and tissue types (see **Methods**). Like ChromHMM^7,8^, a widely adopted functional prediction approach, CoRE-BED utilizes binarized data (noting the presence or absence) rather than raw signal of each respective feature. The authors of ChromHMM have demonstrated that utilizing binary data for model training both reduces the risk of overfitting to noise in the raw signal and improves the discovery of regulatory states characterized by chromatin marks with different signal patterns.

Since the upper size limit of a typical CRE is 1 kb^12,13^, we segmented the entire human genome into approximately 3 million 1 kb bins and assessed each bin for overlap with the 19 binarized epigenomic features. We then applied k-modes clustering^14,15^ in conjunction with the elbow method^16^ to learn the optimal number of clusters representing these putative regulatory elements. We found that the data were best explained by nine underlying clusters (Supplementary Figure 1a). Consequently, k-modes clustering using this learned k-value was again performed on an independent dataset – hereafter, the “training” dataset for the decision tree implementation (Supplementary Figure 1b). The model learned the underlying epigenomic features most highly determinate of each of the 9 regulatory classes in the training dataset. Classification performance was tested using a random 10% of the data held out as an independent test set.

The resulting optimal decision tree structure implemented a maximum depth of 15 and achieved a classification accuracy of 0.88 in the test set (Supplementary Figure 1c and Methods). The presence or absence of cohesion subunit RAD21^17^ emerged as the highest-level feature determinate of regulatory state, followed by the active histone mark^18^ H3K4me3. Because there are multiple paths leading to the same regulatory classification, the distribution of epigenomic signatures within each of the nine regulatory classes can be viewed as a spectrum rather than a discrete class definition, with a different feature distribution for each class (Supplementary Figure 2). Informed by the putative functions of epigenomic signatures as described in the existing literature, we assigned a functional label to each of the nine learned categories.

We term Class 0 “non-functional” due to these regions’ lack of strong enrichment for features associated with highly active chromatin, as well as their relatively uniform distribution across tissues (Supplementary Figure 3a). We call members of Class 1 “distal insulators” due to their enrichment for CTCF and cohesion subunits SMC3 and RAD21^19^, along with their distance from a gene transcription start site (TSS). Class 2 is termed “active promoter 1,” as it is characterized by enrichment for H3K4 methylation, proximity to a gene TSS, and chromatin accessibility^20,21^, while Class 3 is “transcribed gene body” because of its enrichment for H3K36me3, H4K20me1, and H3K79me2^22^. Class 4 elements, which we term Development Associated Elements (DAEs), are enriched for either the dual presence of H3K4me2 and H3K9ac, along with proximity to a TSS, or alternatively, H4K20me1 and H3K79me2. These combinations have been previously described individually but never together. Specifically, H3K4me2 and H3K9ac together are associated with kinetochore assembly at the centromeres, suggesting a role in active cell division^23^. H4K20me1 and H3K79me2, on the other hand, are together associated with transcriptional pause release^24^, an essential mechanism of organismal development^25^. Cumulatively, elements of this class thus show enrichment for two development-associated epigenetic signatures, supporting the DAE moniker.

Class 5 elements are “proximal insulators” due to the strong presence of SMC3, CTCF, and RAD21^26,27^, in addition to TSS proximity. We call Class 6 elements “active promoters 2” due to their harboring of H3K4me2 and H3K4me3, as well as TSS proximity^28^. What differentiates these elements from the active promoters in Class 2 is enrichment of bound RAD21. Class 7 elements are “active enhancers” due to their strong enrichment for H3K27ac, H3K4me1, and accessible chromatin regions^29,30^. Finally, Class 8 appears to capture a distinct category of enhancer-like promoters, hereafter referred to as ePromoters. Elements in this class are enriched for both signatures of active enhancers (acetyltransferase EP300 and H3K27ac)^31^ and active promoters (H3K4me3, H3K9ac, and POL2RA)^28,32,33^.

The distribution of each learned functional category (excluding the “non-functional” Class 0) was variable across each of the 33 cell and tissue types. Class 1 (distal insulator) was most highly enriched in mesenchymal stem cells, Class 2 (active promoter 1) in the peripheral nervous system, Class 3 (transcribed gene body) in embryonic stem cells, Class 4 (DAE) in ESC-derived cell types and bone, Class 5 (proximal insulator) and Class 6 (active promoter 2) in cancer, Class 7 (active enhancer) in embryonic stem cells, and Class 8 (ePromoter) in bone (Supplementary Figure 3).

Using the optimal CoRE-BED decision tree, we generated genome-wide functional annotations on 816 EpiMap^34^ biosamples encompassing 33 distinct cell and tissue types (adipose, blood and T-cell, bone, brain, cancer, digestive, endocrine, endothelial, epithelial, embryonic stem cell-derived, embryonic stem cell, eye, heart, hematopoietic stem cell (HSC) and B-cell, induced pluripotent stem cell, kidney, liver, lung, lymphoblastoid, mesenchyme, muscle, myosatellite, neurosphere, pancreas, placenta and EMM, peripheral nervous system, reproductive, smooth muscle, spleen, stromal, thymus, and urinary). When hierarchically clustering all 816 biosamples based on similarity of genome-wide CoRE-BED functional annotations across the same loci, we observed that biosamples clustered together based on tissue of origin (Figure 2a). Given the 19 epigenomic features, binarized as present or absent, we would expect a total of 2^19^ = 524,288 possible feature combinations across the dataset. However, only a fraction of these expected combinations (35,991) is actually observed across all 816 biosamples. This small core set of epigenomic combinations relative to the number of possible combinations suggests that similar to DNA mutation, epigenomic marks are also under functional constraint^35,36^. Supporting this hypothesis is the observation that cancer cells, characterized by high level of epigenomic dysregulation^37^, harbor the highest number of unique combinations of epigenomic marks in putative regulatory elements across all tissues (3,365). This is nearly three times the number of unique epigenomic signatures shared across all 33 cell type and tissue categories (1,214) (Figure 2b).

**Figure 2.**
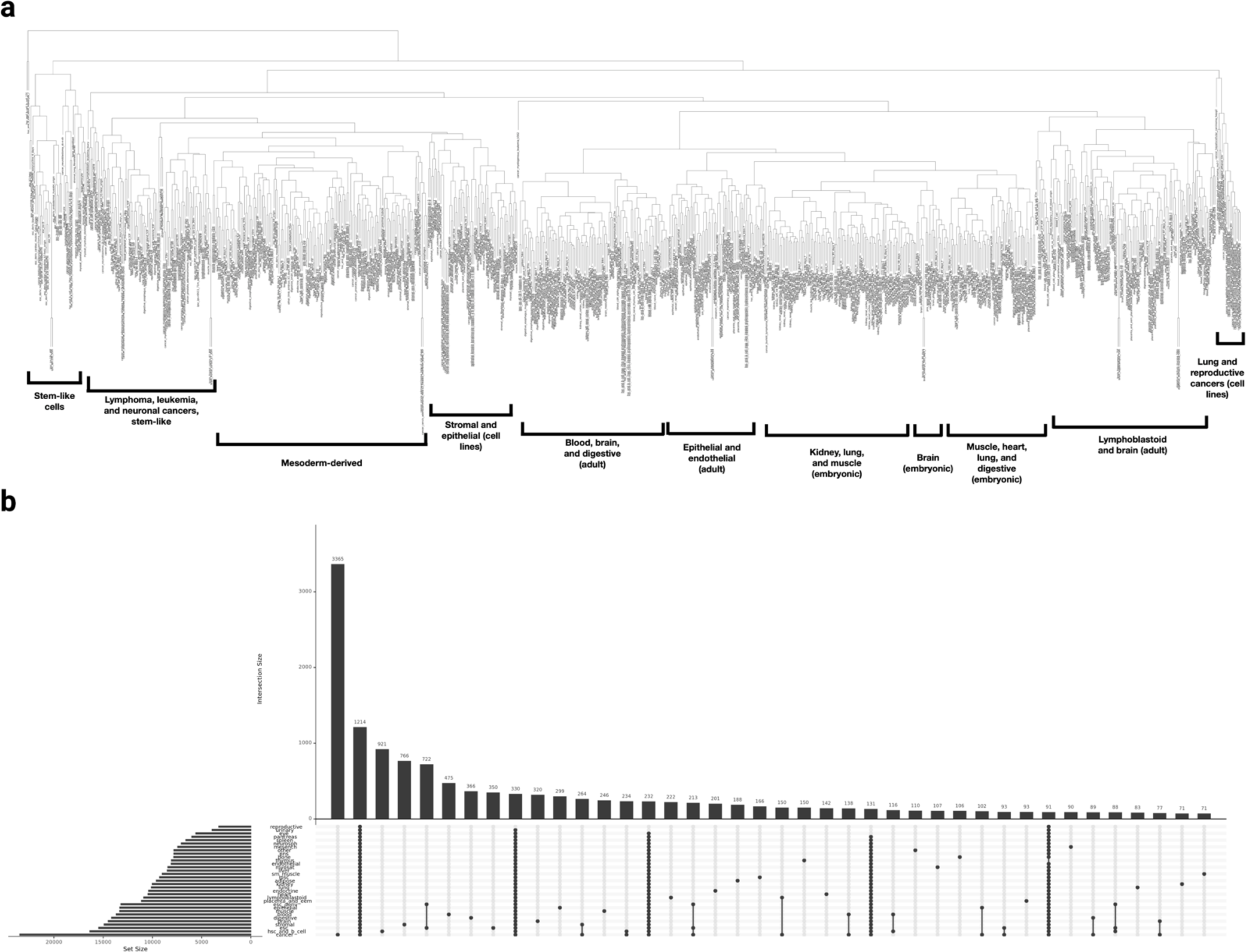
The distribution patterns of CoRE-BED functional classes are non-random. **a,** Hierarchical clustering of 816 biospecimens based on genome-wide functional annotations at 1 kb resolution. Clustering reveals discrete groupings of samples by cell potency and differentiation pathways. **b,** Across all 816 biosamples, observed combinations of the 19 epigenomic features utilized by CoRE-BED were aggregated together by cell type or tissue of origin. The resulting UpSet plot depicts how these unique combinations were shared across all 33 cell and tissue categories. Of the 524,288 (2^19^) possible feature combinations, only 35,991 were observed across the 33 cell and tissue types. 1,214 were shared across all tissue categories, while the largest number of these feature combinations (3,365) was unique to cancer, suggesting disease-related epigenomic dysregulation.

### Comparison of CoRE-BED regulatory annotations with ENCODE cCREs

The CoRE-BED catalog of functional annotations presented here is not the first tissue-wide resource of functional elements. The ENCODE Consortium, for example, has published a registry of 1,063,878 cCREs across 1,518 cell types^5^. In contrast to the learned decision tree-based approach of CoRE-BED, the ENCODE approach^5^ classifies cCREs into one of five pre-defined functional categories (Promoter-like, Proximal enhancer-like, Distal enhancer-like, CTCF-only, and DNase-H3K4me3). These categories are based on the presence of pre-defined biochemical signatures, which can be applied universally to a cell or tissue type and were not learned agnostically.

Because of the high overlap between CoRE-BED’s functional categories and those defined in the ENCODE cCRE registry, ENCODE cCREs present a useful dataset with which CoRE-BED’s regulatory annotations can be more systematically compared. Because the size of the functional annotations is inherently different between the two datasets (uniform 1 kb bins for CoRE-BED versus a variable window size based on DHS signal for ENCODE cCRE), overlapping regions (by at least 1 bp) classified as functional in any cell or tissue type by CoRE-BED and the cCRE registry, respectively, were first merged into single regions. This resulted in a set of 290,846 CoRE-BED functional elements and 1,057,868 ENCODE cCREs. Notably, 186,544 of the global CoRE-BED functional regions (64.14%) were also classified as functional in the cCRE registry, while the remaining 104,302 (35.86%) were unique to CoRE-BED (Jaccard similarity index = 0.19). Among these CoRE-BED-specific annotated regions, most appeared to be Class 3 (transcribed gene body) elements, a category not included in the ENCODE cCRE registry. ENCODE cCRE registry also measured the genomic footprint of these predicted functional elements, estimating that their compendium of regulatory regions covers 7.9% of the entire genome. We quantified the coverage for our tissue-wide CoRE-BED predictions and estimated that they cover 23.72% of the entire genome.

### Comparing CoRE-BED functional classes with ChromHMM 18-state model

Next, we compared CoRE-BED’s tissue-wide regulatory element predictions across the eight functional classes (excluding the non-functional Class 0) with those generated using ChromHMM’s 18-state Roadmap model in the same underlying EpiMap samples^34^. Unlike ENCODE cCREs, which are called using a set of pre-defined epigenetic signatures, ChromHMM utilizes a multivariate hidden Markov model trained on six histone marks (H3K4me3, H3K4me1, H3K36me3, H3K27me3, H3K9me3, and H3K27ac) to annotate genomic regions as belonging to one of 18 learned chromatin states^7,8^. Because the CoRE-BED model was trained using many of the same underlying EpiMap samples as the 18-state ChromHMM Roadmap model, we investigated their degree of sharing of functional categories.

Tissue-wide functional predictions generated from EpiMap data were compiled for each of the eight CoRE-BED functional classes (excluding Class 0) and each of the 18 ChromHMM chromatin states, respectively. Genomic regions overlapping by at least 1 bp in the resulting sets were merged. Next, we computed pairwise Jaccard similarity coefficients to assess how analogous each CoRE-BED functional class was to each of the chromatin states learned by ChromHMM. While Classes 1 (distal insulator), 3 (transcribed gene body), 5 (proximal insulator), 7 (active enhancer), and 8 (ePromoter) from CoRE-BED had a Jaccard index of 0.18 or greater with at least one ChromHMM chromatin state (Supplementary Figure 4), Classes 2 (active promoter 1), 4 (DAE), and 6 (active promoter 2) showed little to no overlap with any of the 18 chromatin states learned by ChromHMM. This suggests that despite encompassing fewer regulatory categories, the CoRE-BED model captures specific patterns of chromatin marks that evade detection by ChromHMM. Each of ChromHMM’s 18 chromatin states, however, had a Jaccard index of 0.14 or above with at least one of the eight CoRE-BED functional classes.

Among the five CoRE-BED functional classes that did show high overlap with ChromHMM chromatin states, we found that each CoRE-BED class and its corresponding most similar ChromHMM chromatin state displayed shared biological properties. For example, CoRE-BED’s Class 1 (distal insulator) overlapped most highly with ChromHMM’s EnhA1 (Active Enhancer 1) (Jaccard = 0.38), both of which are distal elements.

Because CoRE-BED is trained using CTCF and components of the cohesion complex, however, it is able to distinguish between enhancers and insulators, while ChromHMM’s 18-state model does not classify insulators. CoRE-BED’s Class 3 elements (transcribed gene body) showed the strongest similarity to ChromHMM’s ZNF/Rpts (ZNF genes and repeats) (Jaccard = 0.30), both of which identify genes. CoRE-BED’s Class 5 (proximal insulator) was most similar to ChromHMM’s TssA (Active TSS) (Jaccard = 0.18), both of which are proximal to a gene. Finally, CoRE-BED’s Class 7 (active enhancer) showed highest overlap with ChromHMM’s TssFlnkU (Flanking TSS Upstream) (Jaccard = 0.37), while Class 8 (ePromoter) showed highest similarity with ChromHMM’s TssA (Active TSS) (Jaccard = 0.34).

### Characterizing gene expression under CoRE-BED active promoter control in the peripheral nervous system

One CoRE-BED regulatory category that showed low overlap with ChromHMM’s 18 chromatin states was Class 2 elements (active promoter 1), for which the highest Jaccard similarity coefficient across all ChromHMM chromatin states was only 0.03 (with TssA). If the Class 2 elements are indeed a type of active promoter, then one should expect them to be proximal to highly expressed genes. Of the 33 CoRE-BED cell and tissue types, we found that the peripheral nervous system (PNS) was most highly enriched for Class 2 elements (Supplementary Figure 3c). Therefore, we investigated the expression of genes proximal to PNS-specific active promoters using tibial nerve expression data from 619 individuals in GTEx. Of the 1,976 genes proximal to a Class 2 CoRE-BED active promoter in the PNS, 1,730 (87.55%) were expressed in tibial nerve.

Among these 1,730 genes, the median expression was substantially higher than that of the remaining 53,755 expressed genes in tibial nerve (median TPM of genes proximal to Class 2 elements: 3,172.315, median TPM of remaining expressed genes: 1.947, p < 2.2 x 10^-16^), supporting the “active promoter” nomenclature.

### Characterization of enhancer-like promoters

Another distinctive regulatory class learned by CoRE-BED is the enhancer-like promoter, or ePromoter (Class 8), a previously-described class characterized by both promoter-like and enhancer-like activity^38–40^. Notably, among the ePromoters, we observed both a higher proportional overlap with enhancer RNAs (0.07 versus 0.02) and a shorter median distance (22.02 kb versus 64.80 kb; p < 2.2 x 10^-16^) from a transcribed enhancer RNA relative to all other putative regulatory elements across the genome. Additionally, we found an enrichment for biological pathways associated with general cell function and maintenance of homeostasis in genes proximal to ePromoters, consistent with previous studies linking promoters with enhancer activity to housekeeping genes^38^ (Figure 3a). Finally, we observed highly significant enrichment (Benjamini-Hochberg-corrected p < 0.05) of 192 transcription factor binding motifs (Supplementary Table 1) that were unique to ePromoters (see **Methods**). Among these motifs were AP1 and members of the STAT and ATF/CREB families, whose enrichment has previously been associated with enhancer-like promoters^38^.

**Figure 3.**
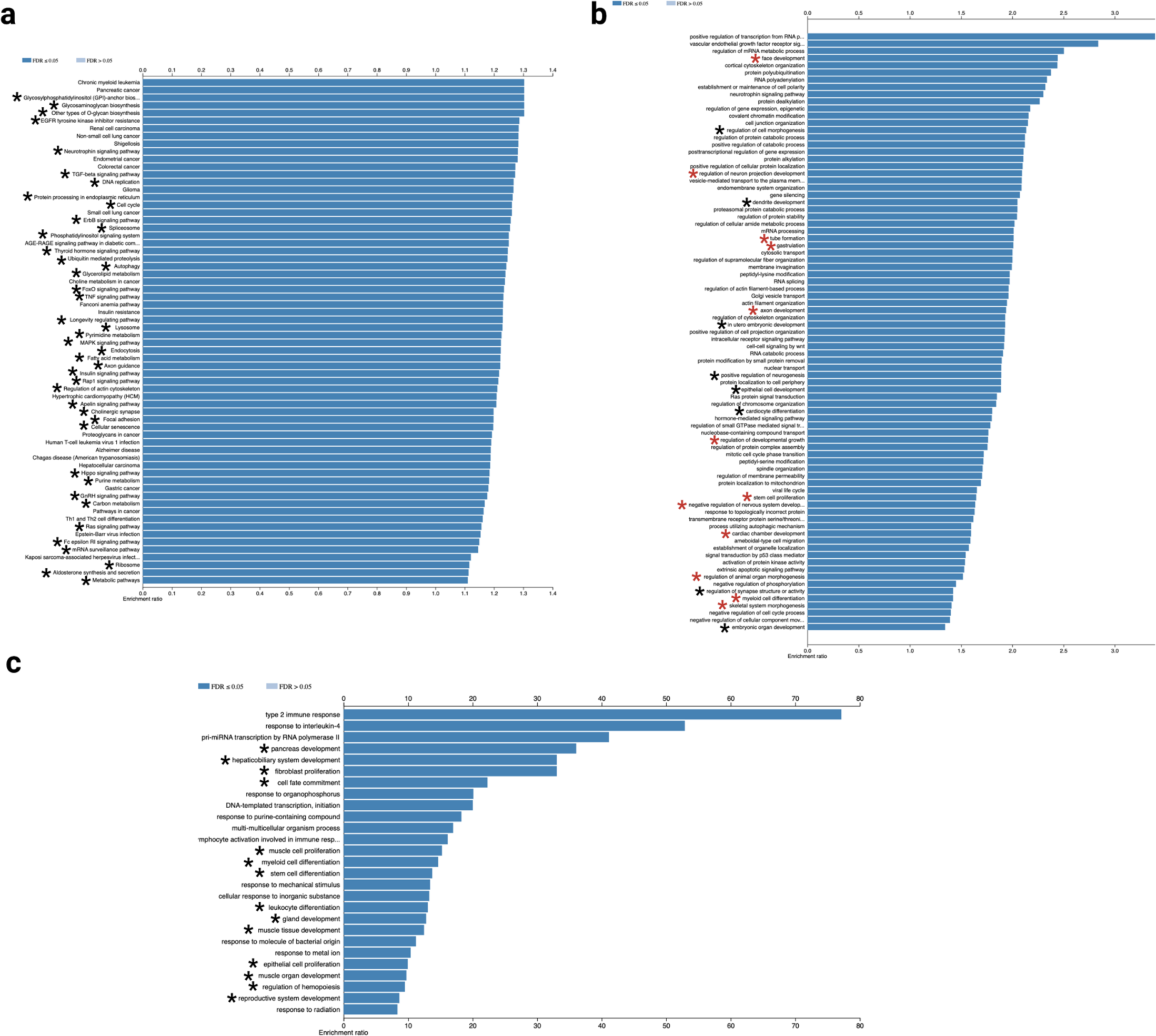
CoRE-BED ePromoters and DAEs are non-randomly distributed and enriched for gene pathways associated with development. **a,** Genes proximal to an ePromoter in at least one cell or tissue type were found to be enriched for housekeeping pathways associated with general cell function and maintenance of homeostasis (marked with asterisk). This observation coincides with previous reports of enhancer-like promoters. **b,** Genes proximal to a DAE in ESCs (marked simultaneously by H3K4me2 and H3K9ac) are enriched for pathways associated with embryonic development (marked with asterisk). Pathways marked with a red asterisk are enriched ESC pathways that were not significantly enriched in thymus, the tissue with the lowest proportion of DAEs. **c,** Gene ontology terms over-represented among transcription factors with binding sites at DAEs. Terms associated with developmental pathways are marked by an asterisk.

### CoRE-BED identifies class of DAEs enriched in stem-like cell types

CoRE-BED identified a previously uncharacterized class of regulatory elements (Class 4), i.e., the DAEs, a unique chromatin state defined by two distinct epigenomic feature combinations (either H3K4me2 and H3K9ac or H4K20me1 and H3K79me2). While these combinations have previously been described individually, with H3K4me2 and H3K9ac associated with kinetochore assembly^23^ and H4K20me1 and H3K79me2 with transcriptional pause release^24^, they have never been viewed as defining a single regulatory category. Because each pair of histone marks is associated with a key developmental process, we termed this class “development-associated elements,” or DAEs.

Supporting the DAE nomenclature was these elements’ enrichment in stem-like cell types. Across each of the 33 major cell and tissue types, we calculated the mean number of CoRE-BED DAEs. In concordance with what one would expect for development-associated elements, three of the top four most highly enriched cell types were embryonic stem cell (ESC)-derived cells (mean 687.87 DAEs per biosample), ESCs (mean 500.83 DAEs per biosample), and myosatellite cells (mean 444.67 DAEs per biosample). The only non-stem cell type among these most highly enriched tissues was bone, which, however, retains the ability to regenerate in response to a fracture^41^. By comparison, thymus, a terminally differentiated tissue with a generally low regenerative capacity^42^, had the lowest enrichment of DAEs (mean 56.82 DAEs per biosample) (Supplementary Table 2).

Given that CoRE-BED DAEs are highly enriched in stem cells, we next aimed to characterize the function of genes under the putative control of a DAE in ESCs. Among the 2,690 unique genes proximal to a CoRE-BED DAE in one of 8 ESC lines, we ran an over-representation analysis to test for enrichment for KEGG biological pathways^43,44^. Of the 83 KEGG pathways enriched in these genes in ESCs (false discovery rate [FDR] < 0.05), we observed a number of pathways related to embryonic development that were not among the pathways associated with DAE-proximal genes in thymus, the lowest-enriched tissue for DAEs. Among these developmental pathways enriched for in DAEs are face development, regulation of neuron projectin development, tube formation, gastrulation, axon development, regulation of developmental growth, stem cell proliferation, negative regulation of nervous system development, cardiac chamber development, regulation of animal organ morphogenesis, myeloid cell differentiation, and skeletal system morphogenesis (Figure 3b).

A similar pattern was observed for the transcription factors (TFs) predicted to bind at DAEs. Across all 833 unique biosamples analyzed in aggregate, we identified a set of binding sites for 26 TFs enriched at DAEs. Using a subsequent gene over-representation analysis, we found these TFs to be highly enriched for 14 developmental pathways, including pancreas development, cell fate commitment, and stem cell differentiation (Figure 3c). We also subsequently characterized the tissue-specificity of genes under putative DAE control. We obtained tau indices computed across all human genes using GTEx expression data^45,46^, with a tau index of 0 representing ubiquitously-expressed genes and an index of 1 representing tissue-specific genes^47^. We found that genes under the putative control of a DAE had mean tau index of 0.373, representing low-to-moderate tissue-specificity.

Finally, we aimed to characterize how the DAEs resolve during stem cell differentiation. Over the course of embryonic development, for example, bivalent promoters represent a well-characterized “poised” regulatory state in highly pluripotent cell types that is marked by both the active H3K4me3 mark and repressive H3K27me3^48,49^. These bivalent promoters are known to transition over the course of development to either an active or permanently silenced state by losing either H3K27me3 or H3K4me3, respectively. We asked whether the DAEs identified by CoRE-BED transition to or from other regulatory classes in a similar manner. Unlike bivalent promoters, however, we found that most DAEs gained or lost during stem cell differentiation transition directly to or from Class 0 (non-functional) elements. For example, we observed an initial total of 911 DAEs in H1 ESCs, which decreased to 405 in H1 ESC-derived neural crest cells. Of the 684 DAEs lost through differentiation, 598 (87.4%) transitioned to Class 0 (non-functional) elements.

Unlike H1 ESC differentiation to neural crest cells, H9 ESCs instead gained a substantial number of DAEs upon transitioning to smooth muscle cells, increasing from 176 to 3,092 DAEs. Of the 2,976 DAEs gained, 2,907 (97.7%) were classified as Class 0 (non-functional) in non-differentiated H9 ESCs. Generally, DAE loss during ESC differentiation appears to be the result of H4k20me1 depletion, suggesting loss of open chromatin conformation^50^. Conversely, DAEs are gained during ESC differentiation via acquisition of either H3K79me2 or H4k20me1. Because DAEs primarily transition directly to or from a non-functional (Class 0) state, it is unlikely that they represent a transitional class of elements like bivalent promoters. Instead, they represent a discrete class of functional elements.

### A putative role for DAEs in common disease

We explored the impact of DAEs on the genetic architecture of common disease. Across 146,337 SNP associations reported within the NHGRI-EBI GWAS Catalog^51^, 1,697 variants (1.16%) were found to fall within a DAE in at least one cell or tissue type (Supplementary Table 3). Among these associations is the Parkinson’s disease-associated variant rs823128^52,53^, which CoRE-BED classified as a DAE in astrocytes based on the presence of H3K79me2 and H4K20me1 in its immediate vicinity. Interestingly, we also observed enrichment of both H3K9ac and H3K4me2, the other major biochemical signature of DAEs, slightly further upstream of the variant locus (Figure 4).

**Figure 4.**
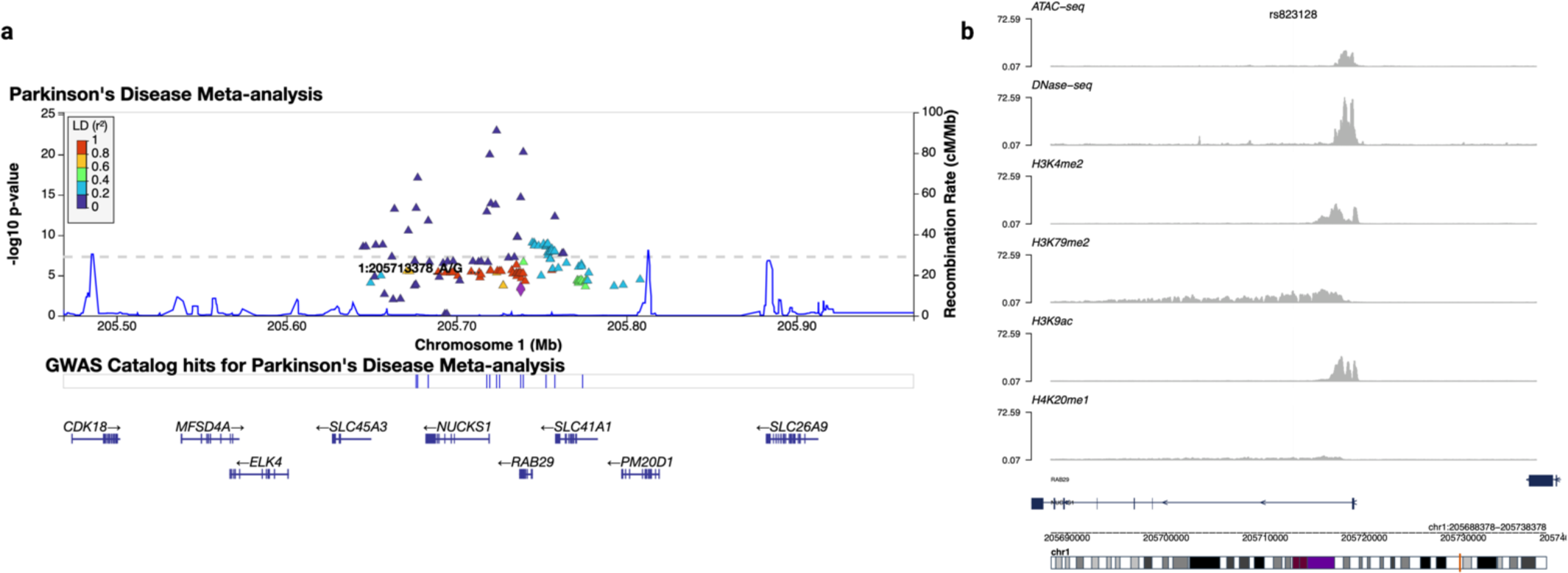
CoRE-BED DAEs overlap with disease-relevant variants. **a,** We identified rs823128, a Parkinson’s disease-associated SNP reported in the GWAS Catalog and replicated in subsequent meta-analyses. This SNP falls within a DAE in astrocytes. **b,** While this SNP region overlaps a DAE based on H3K79me2 and H4K20me1 peaks called in its vicinity, there is also notable enrichment of H3K4me2 and H3K9ac, the second never-before described biochemical signature of DAEs, further upstream near the TSS of NUCKS1.

### DAEs within brain astrocyte eQTLs

Among the top tissues enriched for DAE-associated GWAS SNPs was brain. Of the 375 GWAS Catalog SNPs predicted by CoRE-BED to fall in a DAE in at least one brain cell type, 286 were in astrocytes specifically. Thus, astrocytes appeared to be an ideal model tissue in which to further explore the regulatory role of DAEs in human disease. Using individual-level germline genotyping and gene expression data from human induced pluripotent stem cell (iPSC)-derived astrocytes^54^, we performed expression quantitative trait locus (eQTL) mapping. We identified 126 astrocyte eQTLs within an astrocyte-specific DAE putatively regulating 86 unique genes (Supplementary Table 4). Among these astrocyte eQTLs in a DAE is rs9408928, a SNP associated with the expression *NDUFA8*, a gene previously associated with both Alzheimer’s disease and Parkinson’s disease^55,56^. This SNP has only one immune-related phenotypic association (immune response to smallpox) reported in the NHGRI-EBI GWAS Catalog^51,57^, although no associations with brain-related phenotypes have yet been reported. Within this same set of DAE-overlapping astrocyte eQTLs, we also identified 7 SNPs (rs807748, rs12119677, rs57156794, rs12119878, rs72836878, rs17732024, and rs150792202) associated with the expression of four anxiety-associated genes (*ARVCF*^58^*, FLII*^44,59^*, HMBS*^60^, and *SLC45A1*^61^). None of these SNPs have any phenotypic associations reported in the GWAS Catalog and thus could represent attractive novel targets for future functional experiments characterizing the role of DAEs in disease biology.

### Practical application to GWAS variant prioritization

In addition to providing a genome-wide compendium of regulatory annotations, we hypothesized that the CoRE-BED framework could serve as a powerful tool for improving the functional interpretation of GWAS variants and identifying classes of SNPs that contribute to the heritability of a given trait. The underlying assumption of this hypothesis was that SNPs within a functional region of a phenotypically relevant cell or tissue type should be more likely to causally affect the phenotype of interest. Utilizing a set of SNP associations with at least suggestive significance (GWAS *p* < 1 x 10^-6^) reported across all case-control studies in the NHGRI-EBI GWAS Catalog^51^, we tested whether there was clear, observable stratification between SNPs overlapping a CoRE-BED regulatory element and the remaining SNPs. Among the 25,643 GWAS Catalog SNPs (from 1,613 case-control studies), 11,760 (45.86%) had a CoRE-BED functional annotation in at least one cell or tissue type. As a negative control, we generated CoRE-BED annotations for an equally-sized set of variants, matched for minor allele frequency (MAF), that were sampled from the over 80 million variants cataloged in by the 1000 Genomes Project^62^. In this case, only 6,933 (27.03%) overlapped with a CoRE-BED functional region.

We observed a significantly higher MAF in GWAS Catalog SNPs with a CoRE-BED annotation than those without (with functional annotation median MAF 0.26; without functional annotation median MAF 0.24; p = 4.476 x 10^-13^). We found that the GWAS Catalog variants with a CoRE-BED functional annotation had a more highly significant GWAS p-value distribution than those without (with functional annotation median GWAS p-value 6.99 x 10^-9^; without functional annotation median GWAS p-value 2 x 10^-8^; p-value < 2.2 x 10^-16^), suggesting that genomic variants overlapping with a CoRE-BED functional region may be more highly enriched for causal effects. To test this hypothesis, we obtained a set of fine-mapped SNPs compiled across more than 13,000 GWAS by CAUSALdb^63^. Among the GWAS Catalog SNPs annotated by CoRE-BED, we found that those with a CoRE-BED functional annotation in at least one cell or tissue type showed significantly higher overlap with the CAUSALdb credible SNP set than those without a CoRE-BED functional annotation (55.29% versus 50.36%, two-sample proportion test *p* = 9.51 x 10^-16^).

Among the 11,760 GWAS Catalog SNPs from a case-control study with a CoRE-BED annotation in at least one cell or tissue type, the highest proportion (52.41%) were annotated as distal insulator variants in at least one cell type, followed by transcribed gene body variants (potentially involved in alternative transcript splicing) (52.14%), active enhancer (27.52%), ePromoter (21.84%), proximal insulator (19.48%), active promoter (type 2) (3.69%), DAEs (2.30%), and active promoter (type 1) (0.93%). Notably, a high proportion (73.80%) of CoRE-BED-annotated variants also overlapped with a fine-mapped cis-eQTL identified by GTEx.

To illustrate a CoRE-BED application to GWAS, we utilized CoRE-BED to predict functional status of SNPs from a GWAS meta-analysis of hypertension from the U.K. Biobank^64^. Because this was a cardiovascular phenotype, functional annotation was limited to heart tissues. Of the 28,987,534 variants included in the GWAS, 1,487,383 (5.13%) overlapped with a CoRE-BED functional element in heart.

Comparing overall p-value distributions between SNPs with a CoRE-BED functional annotation in heart versus all remaining GWAS variants, we found a significantly lower GWAS p-value distribution among those with a CoRE-BED functional annotation (median p-value of 0.4597 versus median p-value of 0.4652; p-value < 2.2 x 10^-16^; Figure 5a)^65^. We then wondered whether these CoRE-BED prioritized variants might also be more highly enriched for phenotype heritability. To test this hypothesis, we utilized partitioned linkage disequilibrium score regression (LDSC) to estimate the proportion of SNP heritability from the hypertension GWAS that is captured by heart-specific CoRE-BED regulatory elements. The broad-sense heritability of hypertension has been estimated to be between 30% and 50%^66,67^. Although only 5.13% of GWAS variants had CoRE-BED annotations in heart tissue (Figure 5b), these CoRE-BED-annotated variants were estimated to capture nearly all of the total SNP heritability (*h*^2^ = 0.045 overall, Figure 5c). In terms of individual regulatory categories, Class 1 (distal insulators) accounts for the largest proportion of SNP heritability captured by the GWAS (49.5%, *h*^2^ = 0.022), followed by Class 7 (active enhancers) (44.2%, *h*^2^ = 0.020).

**Figure 5.**
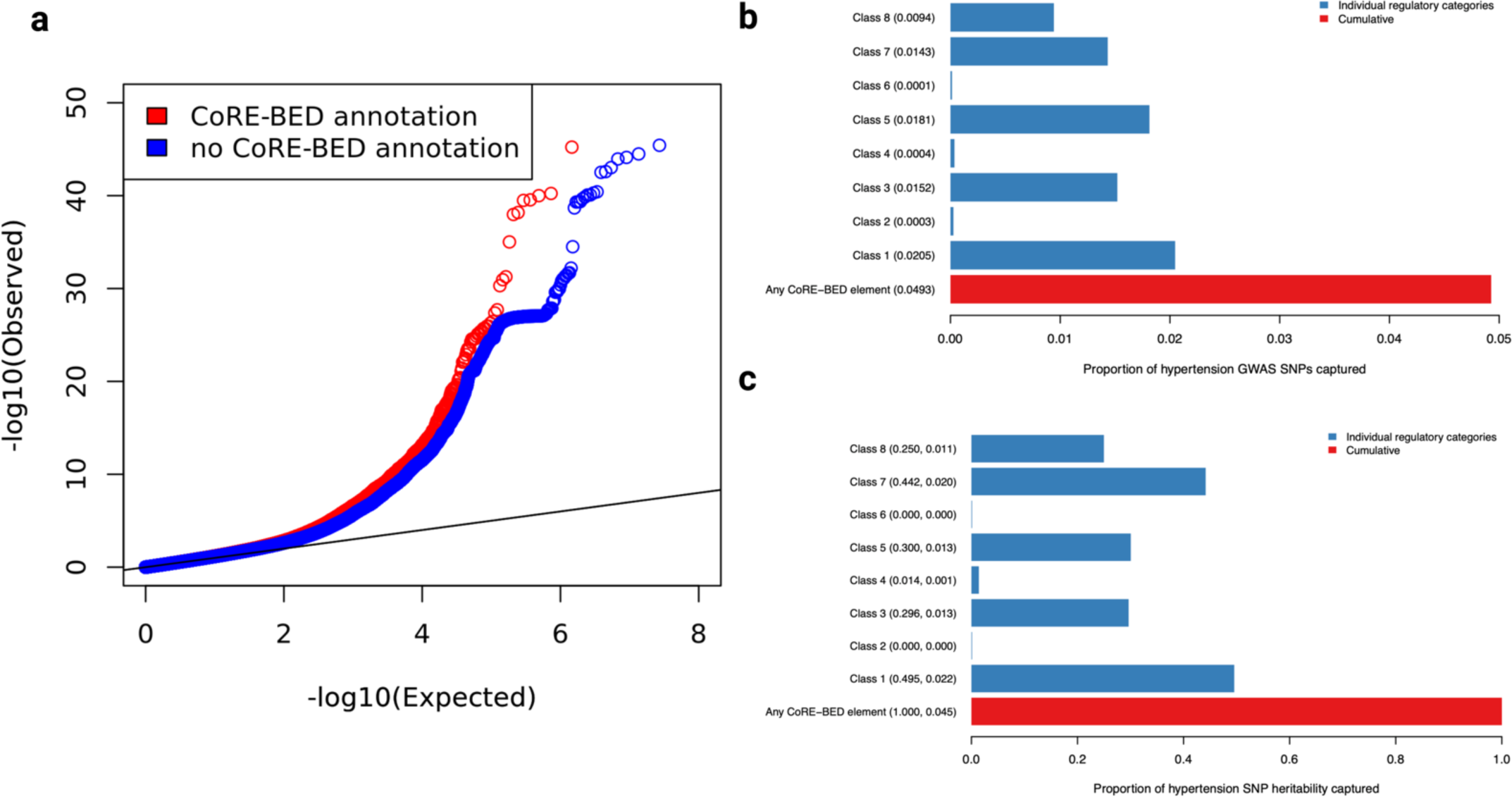
CoRE-BED functional annotations capture the SNP heritability from a GWAS of hypertension. **a,** Across all 28,987,534 variants tested in a GWAS of hypertension, the 1,487,383 (5.13%) with CoRE-BED annotations in heart had a significantly lower p-value distribution than the remaining SNPs. **b,** The proportion of SNPs included in the partitioned heritability estimation with a CoRE-BED annotation for each functional category. **c,** The proportion of SNP heritability explained by each CoRE-BED functional category. Values in parentheses represent the proportion of SNP heritability and the overall SNP heritability, respectively.

### Post-GWAS analysis: phenome-wide heritability by CoRE-BED annotations and causal variant identification

Given CoRE-BED’s effectiveness at capturing SNPs contributing to disease heritability in the hypertension GWAS, we next applied this same methodology^68–70^ more broadly across a set of 69 complex traits^71–73^ (Supplementary Table 5). Strikingly, across these 69 traits, we observed a result similar to the hypertension finding. Interestingly, however, the relative contributions of each CoRE-BED functional class to trait heritability were highly variable across phenotypes (Figure 6a). Across all traits, using LDSC again, Class 1 (distal insulator) elements were predicted to explain the greatest mean proportion (48.75%) of SNP heritability (Supplementary Figure 5b), followed by Class 3 (transcribed gene body) with a mean proportion of 47.95% (Supplementary Figure 5d) and Class 7 (active enhancer) with a mean proportion of 45.64% (Supplementary Figure 5h).

**Figure 6.**
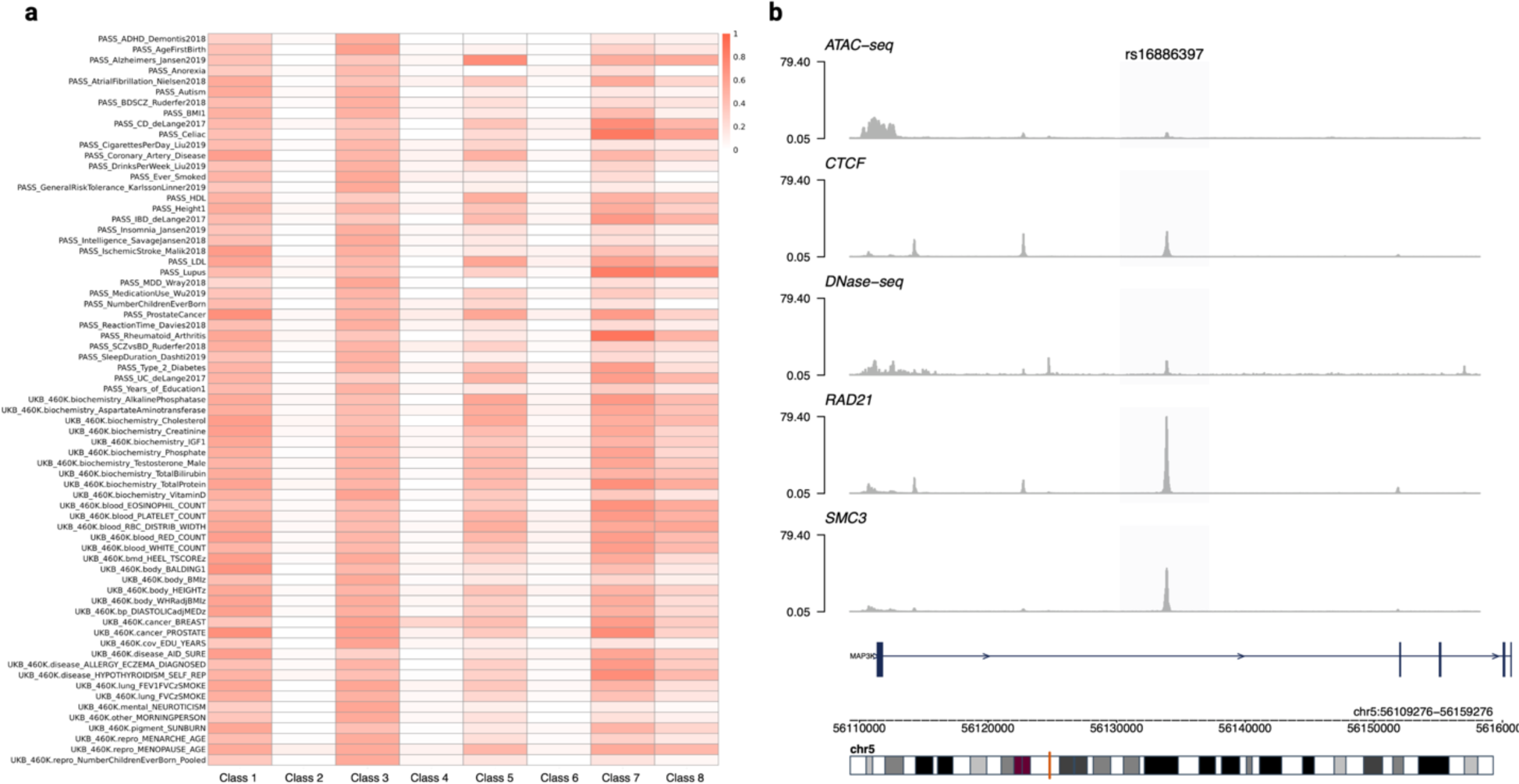
Applicability of CoRE-BED to post-GWAS analysis. **a,** We estimated the proportion of SNP heritability attributable to CoRE-BED regulatory predictions using GWAS of 69 complex traits performed in up to 1.03 million individuals. Relative contributions varied by trait, with Class 1 (distal insulators) showing the greatest mean contribution across all traits. **b,** CoRE-BED accurately predicts function in gold standard list of causal variants. The breast cancer-associated SNP rs16886397 was functionally validated to confirm cS2G’s prediction of MAP3K1 as the target gene. CoRE-BED predicts this SNP to be within a proximal insulator (Class 5 element). We observed an enrichment of ChIP-seq signal for CTCF, RAD21, and SMC3 at the SNP locus, all known signatures of insulator elements. Additionally, ATAC-seq and DNase-seq signal enrichment was present at the same locus, indicating open chromatin at the site. The SNP localizes to an intron of MAP3K1, supporting CoRE-BED’s classification of a proximal versus distal insulator and demonstrating CoRE-BED’s potential usefulness in identifying targets for functional studies.

Finally, we leveraged a set of 25 experimentally validated causal SNP-gene pairs published as part of a recent study^73^ to assess CoRE-BED’s ability to functionally identify true causal variants. Of the 25 experimentally validated SNPs in this dataset, 17 (68%) overlapped with a CoRE-BED functional element in at least one cell or tissue type. Of these 17 SNPs, 9 were Class 1 elements (distal insulators) in at least one cell or tissue type, 9 were classified as Class 3 (transcribed gene body), 4 as Class 5 (proximal insulators), 7 as Class 7 (active enhancers), and 4 as Class 8 (ePromoters). Among these validated SNPs was rs16886397, an intronic variant of *MAP3K1* associated with breast cancer risk. Within the MCF 10A breast cancer cell line, CoRE-BED predicted this SNP to be within a proximal insulator. In concordance with this prediction, we observed distinct, overlapping ChIP-seq signals at the SNP locus for CTCF, RAD21, and SMC3, key insulator signatures, as well as DNase-seq and ATAC-seq signals, indicating open chromatin at the variant position (Figure 6b).

In comparison with CoRE-BED, the same number (17) of (non-identical) functionally validated SNPs overlapped an ENCODE cCRE. Cumulatively, 21 of the 25 (84%) SNPs are captured using the two predictive approaches together, demonstrating the potential utility of an ensemble approach for functional prediction. Because ChromHMM assigns states across the entire genome, all 25 of these SNPs overlapped with one of these 18 chromatin states in any given tissue. Among the eight functionally validated SNPs without a CoRE-BED functional annotation, we observed overlap with weak transcription (TxWk), weak enhancer (EnhWk), heterochromatin (Het), repressed Polycomb (ReprPC), weak repressed Polycomb (ReprPCWk), and quiescent (Quies) chromatin states, all of which would be categorized as “non-functional” by CoRE-BED.

## Discussion

We present CoRE-BED, a tree-based framework trained using 19 epigenomic features across 33 major cell and tissue categories, achieving high predictive performance in an independent test set. The major benefit of a decision tree-based architecture is the ease of interpretability and the transparency regarding which underlying features drive each prediction. Using our accompanying command line tool, the CoRE-BED model can be leveraged to annotate a set of genomic coordinates, including full GWAS summary statistics.

Intriguingly, of the 19 epigenomic features, we observed only 35,991 unique combinations of features across the more than 2.5 billion total 1 kb bins considered across the 816 biosamples used to compile the training and test datasets. If these features were distributed randomly throughout the genome, we would expect to observe well over 500,000 unique combinations (2^19^). This discrepancy between possible and observed feature combinations suggests some mechanistic constraint, under which only certain epigenomic feature combinations are compatible with cellular function. Supporting this hypothesis is our finding that a sizable proportion of the total observed feature combinations were unique to cancer cells, which are characterized by widespread epigenomic dysregulation^37^. These 3,365 cancer-specific feature combinations might serve as attractive targets for future functional validation studies, in which any resulting changes in gene expression or cell morphology could be assessed after induction of cancer-specific epigenomic feature patterns.

After training an optimal decision tree classifier, we generated functional annotations across 816 individual biosamples encompassing the 33 major cell and tissue types represented by CoRE-BED^34^. Notably, rather than a rigid set of definitions for each functional class, each CoRE-BED class can be viewed as a discrete distribution of epigenomic features. Under this model, there is no single set of features that define a given regulatory class. However, there is a specific subset of epigenomic features with a higher likelihood for each type of functional element. This dynamic model is in some ways analogous to the degeneracy of the genetic code itself, in which wobble base pairing allows for multiple codons to code for a single amino acid.

An interesting regulatory class identified by CoRE-BED is the class of Development Associated Elements, or DAEs. Enriched in tissues with increased regenerative capacity, these elements are defined by either of two distinct histone mark combinations, either H3K4me2 and H3K9ac or, alternatively, H4K20me1 and H3K79me2. These histone mark combinations have previously been described individually, with H3K4me1 and H3K9ac associated with kinetochore assembly^23^ and H4K20me1 and H3K79me2 with transcriptional pause release^24^. However, this is the first time that both sets of features, each associated with a key development-associated mechanism, have been grouped together to define a regulatory class. Among SNPs reported in the GWAS Catalog that overlap with a CoRE-BED DAE, we observed both broad H3K79me2 and H4K20me1 signal and punctate H3K4me2 and H3K9ac in the vicinity of DAE-associated SNP loci (Figure 4). This suggests that these two histone mark combinations may tend to co-localize to shared regions of the genome, which may in-part explain why the CoRE-BED model considers them as a single regulatory class.

At the outset of training the CoRE-BED model, we utilized k-modes clustering in conjunction with the elbow method to determine the suitable number of clusters characterizing our training data (Supplementary Figure 1). While simple and easy to implement, the use of the elbow method also introduces some inherent amount of subjectivity, as one must visually distinguish what number of clusters appears to be optimal^74^. The choice of a larger number of clusters on which to train the CoRE-BED model, rather than the original nine, may lead to the two distinct sets of chromatin marks that compose DAEs to be further resolved into two separate regulatory categories. However, utilizing too many clusters might dilute important relationships between different chromatin marks, as well as the degeneracy of the regulatory code within an individual regulatory class.

We demonstrate CoRE-BED’s ability to prioritize GWAS variants based on tissue-specific functional annotation, both among genome-wide significant variants reported to the NHGRI-EBI GWAS Catalog and within GWAS summary statistics from 70 complex traits. Notably, distal insulators encompassed the largest proportion of heritability across these traits. Insulators play a key role in modeling three-dimensional genomic architecture and establishing boundaries between topologically associated domains (TADs)^75,76^, and it has been recently reported that these TAD boundaries are enriched for complex trait heritability^77^. The results reported here support this finding and provide a comprehensive reference of insulator contributions to disease heritability, predicting both proximal and distal sub-classes.

In addition to the case-based limitations discussed above, the CoRE-BED implementation has several limitations at the model level. For instance, the classifier from a decision tree structure may be highly dependent on the composition of the training data. To address this issue, averaging techniques, including bagging and random forests, may be used, reducing the problem of high variance. However, given the scale of the epigenome-wide data, these approaches (and related ones) are computationally onerous. For this reason, we tested CoRE-BED in additional epigenomic data. Another limitation is that CoRE-BED’s methodological simplicity may imply loss in power.

As a decision tree, CoRE-BED is scale-invariant. A split on a feature is not influenced by the other features. No feature scaling is required, and the classifier is invariant to monotonic transformations of an epigenomic feature. Decision tree classifiers are, in this regard, inherently different from model-based methods such as neural networks.

The methodological simplicity of CoRE-BED greatly enhances the interpretability of its functional predictions. As demonstrated here, integrating CoRE-BED functional predictions into a genomic analysis workflow enhances understanding of the functional context of regions across the genome. Utilizing the CoRE-BED framework as one pillar of an integrative, multi-method approach of investigating genotype-phenotype relationships opens new avenues for potential study.

## Methods

### CoRE-BED classifier

The CoRE-BED framework is trained using 19 features (TSS proximity; histone marks H3K27ac, H3K27me3, H3K4me1, H3K4me2, H3K4me3, H3K36me3, H3K79me2, H3K9ac, H3K9me3, and H4K20me1; chromatin accessibility assays ATAC-seq and DNase-seq; as well as CTCF, EP300, H2AFZ, POL2RA, RAD21, and SMC3 binding) across 33 major cell and tissue types (adipose, blood and T-cell, bone, brain, cancer, digestive, endocrine, endothelial, epithelial, ESC-derived, ESC, eye, heart, HSC and B-cell, iPSC, kidney, liver, lung, lymphoblastoid, mesenchymal stromal, muscle, myosatellite, neurosphere, other, pancreas, peripheral neuron, placenta, reproductive, smooth muscle, spleen, stromal, thymus, and urinary) (Figure 1). The raw data were originally published as part of the EpiMap repository^34^. Because the average size of a genomic regulatory element (i.e., promoter or enhancer) is approximately 1 kb^78,79^, the entire genome was first segmented into kilobase-resolution, non-overlapping bins. The result was an initial genome-wide set of 3.2 million regions^80^ on which to conduct the training for the classifier. Overlap was calculated between each 1 kb bin and the 19 genomic features^34^ in 33 biosamples representing each of the cell and tissue types.

The resulting training data (presence or absence of each of the 19 genomic features in each 1 kb bin) are unlabeled, meaning there is no starting “ground truth.” Thus, k-modes clustering^14,15^, in conjunction with the elbow method^16^, was applied to the 33 biosamples, resulting in nine clusters of putative *cis*-regulatory elements that best captured the data. K-modes clustering was then repeated using this k = 9 to derive regulatory class labels (Class 0 – Class 8) for an independent dataset. This dataset consisted of 66 biosamples, two for each cell or tissue type.

Using these training data and their corresponding k-modes-derived labels, a decision tree classifier was trained using sci-kit learn^81^. A series of maximum depths ranging from 1 to 19 were tested, with node splits determined based on maximal Shannon entropy reduction^11^. The Shannon entropy *H*(*T*) of the tree *T* quantifies the uncertainty of the tree-based classifier. Given the (disjoint) classes of regulatory elements, *x*_1_, *x*_2_,…, *x_k_*, derived from *T*, with corresponding probability *p*(*x*_1_), *p*(*x*_2_),…, *p*(*x_k_*), the Shannon entropy *H*(*T*) is defined as follows:

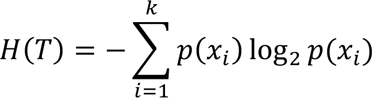

Shannon entropy was a natural criterion because it represents the unique family of functions, defined on the set of all k-dimensional distributions (of the k functional annotations of the classifier), satisfying the Shannon-Khinchin axioms of continuity, symmetry, maximality, and expandability^11^.

A random 10% of the training data was held out to evaluate model performance in an independent test set, with the model using a maximum depth of 15 performing best (88.29% in the test set). Based on which learned combination of genomic features contributed most highly to each class, each of the nine regulatory categories was assigned a name suggestive of putative biological function.

### Generation of CoRE-BED regulatory annotations

The epigenomic features were downloaded from the EpiMap Repository (http://compbio.mit.edu/epimap/) as bigWig tracks based on GRCh37/hg19. The bigWig tracks were converted to bedGraph format before undergoing peak calling using MACS2^82^ to generate final sets of called peaks in UCSC BED format. These resulting peak calls were based on GRCh37/hg19, and an accompanying set based on genome build GRCh38/hg38 was generated using UCSC liftOver^83^. TSS coordinates based on GRCh38/hg38 were downloaded from refTSS^84^, and UCSC liftOver was used to generate a set of TSS coordinates based on GRCh37/hg19. These data were binned at 1 kb resolution and fed into the trained CoRE-BED decision tree to generate genome-wide functional class across all 1 kb bins in 816 unique biosamples.

### Comparison of CoRE-BED functional predictions with ENCODE cCREs

GRCh38/hg38-based cCREs predicted across 1,518 cell types were downloaded from the SCREEN Registry of cCREs V3 (https://screen.encodeproject.org) and lifted over to GRCh37/hg19. For both these ENCODE cCREs and CoRE-BED-predicted functional regions, overlapping coordinates were merged. The degree of sharing between each set of predictions was evaluated using both percent overlap and Jaccard index. We calculated the Jaccard similarity coefficient *s*(*x_i_*, *E*; *T*) of each class *x_i_* of regulatory elements derived from the CoRE-BED decision tree *T* and the set of putative ENCODE cCREs *E* as follows:

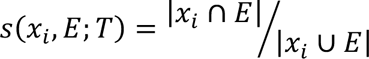

The Jaccard index measures the similarity between the two sets as the proportion of the number of shared elements (i.e., CoRE-BED-predicted regulatory elements in *x_i_* overlapping a putative ENCODE cCRE) relative to the number of distinct elements across the two sets. Note the overall overlap count Ω(*T*, *E*) of the set of *T*-derived regulatory elements with the set of ENCODE cCREs is given by:

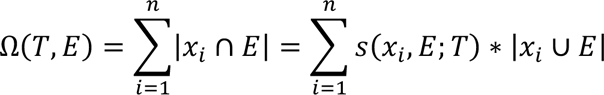

### Comparison of CoRE-BED functional classes with ChromHMM chromatin states

Chromatin state predictions generated across the EpiMap dataset using the ChromHMM 18-state model were downloaded in UCSC BED format and pooled together across all biosamples. These predictions were split up by individual state, and coordinates overlapping by at least 1 bp in these individual datasets were merged. The same process was repeated for the CoRE-BED functional classes, with Class 0 (non-functional) elements being excluded. Using bedtools^85^, we computed pairwise Jaccard similarity indices to determine the degree of similarity of each CoRE-BED functional class with each ChromHMM chromatin state.

### Assessing expression of PNS genes under active promoter control

Class 2 (active promoter 1) CoRE-BED elements were most highly enriched in the PNS. In order to first identify which genes were under the control of these elements in PNS-derived tissues, a list was compiled of all unique genomic regions predicted as Class 2 elements within at least one of 10 PNS-derived biosamples. Using the *closest* function from bedtools^85^, the closest gene to each putative active promoter was identified.

Next, gene-level expression data from tibial nerve, a PNS-derived tissue, was downloaded from the GTEx Consortium^46^ (https://storage.googleapis.com/gtex_analysis_v8/rna_seq_data/gene_reads/gene_reads_2017-06-05_v8_nerve_tibial.gct.gz). Gene expression was quantified using transcripts per million (TPM). Expression levels of genes under the predicted control of a CoRE-BED active promoter were compared with the remaining genes expressed in tibial nerve using a Wilcoxon rank sum test.

### Enhancer RNA (eRNA) enrichment in ePromoters

Class 8 elements learned by the CoRE-BED model were termed ePromoters, promoters with enhancer activity. A comprehensive set of eRNA coordinates was compiled using publicly-available annotation resourced compiled by ENSEMBL^86^, FANTOM5^87^, and Roadmap^88^, in addition to the Human enhancer RNA Atlas (HeRA)^89^. Distance from an eRNA was calculated for each ePromoter and all remaining promoters across the genome using bedtools^85^, and the respective distributions of distances were compared using the Wilcoxon rank sum test^85^.

### Gene pathway over-representation analyses for enhancer-like and DAEs

The closest gene to each CoRE-BED-predicted putative promoter was determined. The resulting sets of genes under putative ePromoter and DAE control, respectively, underwent over-representation analysis using WebGestalt^43^ (http://www.webgestalt.org) to search for enrichment of KEGG^44^ pathway terms. The Organism of Interest was set to “Homo sapiens,” Method of Interest was set to “Over-Representation Analysis (ORA),” and Functional Database was set to “pathway,” with the functional database name set to “KEGG.” The Gene ID Type used was “Gene symbol,” while the Reference Set used was “genome.” Additionally, under “Advanced parameters,” we used an FDR of 0.05 as the significance level.

### Transcription factor binding motif enrichment in enhancer-like promoters and DAEs

The ePromoters and DAEs identified by CoRE-BED underwent separate motif enrichment analysis using the *findMotifsGenome.pl* script from the HOMER suite^90^. A Benjamini-Hochberg p-value correction was used to determine the significance threshold for motif enrichment.

### Computing tau indices for genes under putative DAE control

Tau scores calculated using the GTEx v8^46^ dataset were downloaded from https://genomics.senescence.info/gene_expression/tau.html^45^. The *closest* function from bedtools^85^ was used to determine which GENCODE^91^ gene was closest to each DAE across all 833 tested biosamples, and the corresponding tau index for each of unique gene was retrieved from the reference dataset.

### Predicting GWAS Catalog variants in a DAE

All associations (v1.0) from the NHGRI-EBI GWAS Catalog^51^ (https://www.ebi.ac.uk/gwas/docs/file-downloads) were downloaded, and all reported variants missing genomic coordinates, allele frequency, effect size, or GWAS p-value were filtered out. For the 146,338 remaining SNPs, CoRE-BED functional prediction was performed across all 816 supported biosamples. 1,697 were predicted to be within a DAE in at least one cell type, and 375 of these were specifically predicted to be within a DAE in a brain-derived cell type (286 in astrocytes).

### LocusZoom plot for DAE locus

Once the Parkinson’s disease-associated SNP rs16886397 was identified as a variant of interest, we located the GWAS summary statistics from a recent highly powered meta-analysis that replicated the association. The summary statistics were uploaded to the LocusZoom^92^ web server (http://locuszoom.org) to generate the plot in Figure 4a.

### Trackplot visualizations

The R package trackplot^93^ (https://github.com/PoisonAlien/trackplot) was utilized to visualize bigWig signal at the genomic loci of interest in Figures 4 and 6.

### Describing astrocyte eQTLs within DAEs

Astrocyte data was obtained from the Human Induced Pluripotent Stem Cell Initiative (HipSci)^54^. Using individual-level genotyping and gene expression data from astrocytes derived from differentiated iPSCs grown for 52 days, eQTL mapping was performed. Next, astrocyte-specific CoRE-BED functional annotations were generated across all 71,226 predicted eQTLs. For each of the 126 eQTLs falling within a DAE, over-representation analysis (via WebGestalt^43^) using both KEGG^44^ pathway and Human Phenotype Ontology^94^ terms was performed on the set of genes under the putative *cis-*regulatory control of these DAE eQTLs. Using these pathway-based approaches, genes associated with brain-related phenotypes such as Parkinson’s disease, Alzheimer’s disease, and anxiety were identified.

### Tracking resolution of DAEs during embryonic development

Two ESC lines, H1 and H9, as well as two ESC-derived cell types differentiated from each respective ESC (H1-derived neural crest cells and H9-derived smooth muscle cells), were selected to compare differences in CoRE-BED-DAEs (Class 4 elements). For each pair of cells, we assessed the number of DAEs in the ESC line that were lost after differentiation and which functional categories they transitioned to. We also assessed the DAEs that were gained as a result of differentiation.

### Application to GWAS variant prioritization

Using the filtered set of GWAS Catalog variants described above (see **Predicting GWAS Catalog variants in a DAE**), additional filtering was performed to remove associations with continuous traits, retaining only those from case-control studies. CoRE-BED was used to assign functional status across the resulting variant set across all 816 supported biosamples. Annotations were based on genome build GRCh38. After annotation, variants were stratified based on whether each variant fell within a CoRE-BED functional class (Class 1-8) in at least one cell or tissue type. A Wilcoxon rank sum test with continuity correction was used to statistically compare effect size, p-value, and MAF distributions between variant sets with and without a functional annotation. Significant differences were found for both GWAS p-value and MAF between variants with and without a CoRE-BED functional annotation (see Results). We found no statistically significant difference in odds ratio (OR) between GWAS Catalog SNPs with a CoRE-BED functional annotation and those without (with functional annotation median OR 1.1038; without functional annotation OR median 1.1039; p = 0.079).

As a negative control, to test the expected proportion of randomly sampled variants falling in a CoRE-BED regulatory element, we downloaded the entire set of 2,504 sequenced individuals from the 1000 Genomes Project^62^ at http://ftp.1000genomes.ebi.ac.uk/vol1/ftp/release/20130502/. Using the mean MAF across all populations, we sampled 25,640 SNPs, the same number in our GWAS Catalog set, from among the more than 80 million sequenced variants. The sampled variants were matched for MAF, with each sampled 1000 Genomes variant’s MAF being within 0.01 of the corresponding GWAS Catalog SNP.

### Assessing overlap between CoRE-BED-annotated GWAS Catalog SNPs and fine-mapped credible set

CAUSALdb’s (Version 2.0) credible set of putative causal SNPs^63^ fine-mapped across more than 13,000 publicly available GWAS was downloaded from http://www.mulinlab.org/causaldb/index.html. GWAS Catalog SNPs with and without a CoRE-BED functional annotation were compared for proportional overlap with the credible set of fine-mapped SNPs using a two-sample proportion test, defined by the following formula:

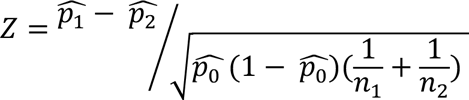

where:

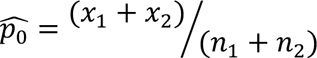

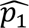 and 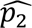 are the observed proportional overlaps of the fine-mapped credible set with GWAS Catalog SNPs annotated and unannotated by CoRE-BED, respectively; *n*_1_ and *n*_2_ are the total number of annotated and unannotated GWAS Catalog SNPs, respectively; and *x*_1_ and *x*_2_ are the number of annotated and unannotated, respectively, GWAS Catalog SNPs overlapping with the fine-mapped credible set.

### Hypertension meta-analysis variant annotation and prioritization

The summary statistics from a multi-ancestry meta-analysis of hypertension conducted in the UK Biobank were downloaded from the Pan-UK Biobank^64^ portal (https://pan-ukb-us-east-1.s3.amazonaws.com/sumstats_flat_files/categorical-20002-both_sexes-1065.tsv.bgz). Functional status was assigned to all GWAS variants, and the summary statistics were subsequently stratified based on the presence or absence of a functional annotation in at least one of 41 heart-derived biosamples. The GWAS p-values of these sets of functionally-stratified SNPs were plotted using the *qqunif* function from the gap R package^95^ and tested for statistically significant divergence using a Wilcoxon rank sum test with continuity correction.

### Partitioned heritability estimation of hypertension and 69 additional GWAS traits

CoRE-BED functional predictions were generated for the 1000 Genomes^62^ European reference set of variants. Annotation status for each functional class was used to generate partitioned linkage disequilibrium scores using LDSC^71,72^. Partitioned heritability based on CoRE-BED functional classes was first estimated for hypertension. Next, for the 69 traits assessed by Gazal et al. to evaluate the performance of cS2G^73^, we leveraged these same partitioned LD scores to estimate the proportion of SNP heritability *h*^2^ captured by variants in each of the 8 CoRE-BED derived functional (Classes 1-8) categories.

## Data Availability

The CoRE-BED model, command line tool, and associated code are publicly available on GitHub (https://github.com/mjbetti/CoRE-BED). Pre-compiled reference files for use with the command line tool, ChIP-seq peak calls generated from EpiMap bigWig tracks, and genome-wide sets of CoRE-BED functional predctions can be retrieved from Zenodo (10.5281/zenodo.7558115).

## Acknowledgment

This research was supported by National Institutes of Health (NIH) grants NHGRI R35HG010718, NHGRI R01HG011138, NIA AG068026, NIGMS R01GM140287, NIMH R01MH126459, and NIH/NCI U01CA253560. Visualizations in Figure 1 were created with Biorender.com.

---

**Extended Figure 1.**
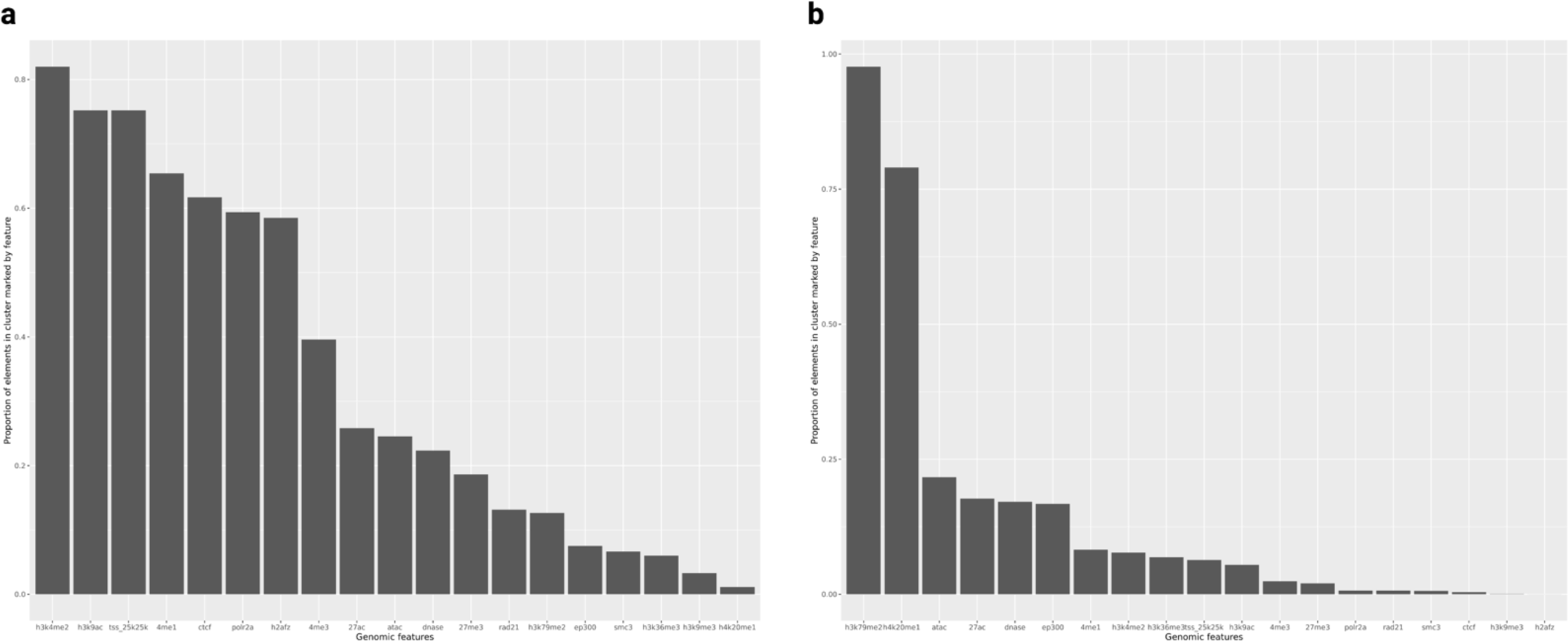
CoRE-BED DAEs are characterized by two previously undescribed combinations of epigenetic markings. **a,** The decision tree training and test sets were composed of 18,522 unique combinations of 19 genomic features. The 783 feature combinations labeled Class 4 (DAEs) in this set were most highly enriched for histone marks H3K4me2, H3K9ac, in addition to TSS proximity. **b,** Across all 173,434 (non-unique) DAE occurrences in 816 individual biosamples representing 33 major cell and tissue types, elements are most highly enriched for H3K79me2 and H4K20me1.

## References

1. Klein, R. J. et al. Complement factor H polymorphism in age-related macular degeneration. Science 308, 385–389 (2005).

2. Cano-Gamez, E. & Trynka, G. From GWAS to Function: Using Functional Genomics to Identify the Mechanisms Underlying Complex Diseases. Front. Genet. 11, 424 (2020).

3. Alsheikh, A. J. et al. The landscape of GWAS validation; systematic review identifying 309 validated non-coding variants across 130 human diseases. BMC Med. Genomics 15, 74 (2022).

4. Soskic, B. et al. Chromatin activity at GWAS loci identifies T cell states driving complex immune diseases. Nat. Genet. 51, 1486–1493 (2019).

5. ENCODE Project Consortium et al. Expanded encyclopaedias of DNA elements in the human and mouse genomes. Nature 583, 699–710 (2020).

6. Fulco, C. P. et al. Activity-by-contact model of enhancer-promoter regulation from thousands of CRISPR perturbations. Nat. Genet. 51, 1664–1669 (2019).

7. Ernst, J. & Kellis, M. Chromatin-state discovery and genome annotation with ChromHMM. Nat. Protoc. 12, 2478–2492 (2017).

8. Ernst, J. & Kellis, M. ChromHMM: automating chromatin-state discovery and characterization. Nat. Methods 9, 215–216 (2012).

9. Murdoch, W. J., Singh, C., Kumbier, K., Abbasi-Asl, R. & Yu, B. Definitions, methods, and applications in interpretable machine learning. Proc. Natl. Acad. Sci. U. S. A. 116, 22071–22080 (2019).

10. Grinsztajn, L., Oyallon, E. & Varoquaux, G. Why do tree-based models still outperform deep learning on tabular data? arXiv [cs.LG] (2022).

11. Khinchin, A. I. & Hinčin, A. J. Mathematical Foundations of Information Theory. (Dover Publications, 1957).

12. Le, N. Q. K., Yapp, E. K. Y., Nagasundaram, N. & Yeh, H.-Y. Classifying Promoters by Interpreting the Hidden Information of DNA Sequences via Deep Learning and Combination of Continuous FastText N-Grams. Front Bioeng Biotechnol 7, 305 (2019).

13. Snetkova, V. & Skok, J. A. Enhancer talk. Epigenomics 10, 483–498 (2018).

14. Huang, Z. & Ng, M. K. A Note on K-modes Clustering. J. Classification 20, 257–261 (2003).

15. Huang, Z. Extensions to the k-Means Algorithm for Clustering Large Data Sets with Categorical Values. Data Min. Knowl. Discov. 2, 283–304 (1998).

16. Marutho, D., Hendra Handaka, S., Wijaya, E. & Muljono. The Determination of Cluster Number at k-Mean Using Elbow Method and Purity Evaluation on Headline News. in 2018 International Seminar on Application for Technology of Information and Communication 533–538 (2018).

17. Cheng, H., Zhang, N. & Pati, D. Cohesin subunit RAD21: From biology to disease. Gene 758, 144966 (2020).

18. Beacon, T. H. et al. The dynamic broad epigenetic (H3K4me3, H3K27ac) domain as a mark of essential genes. Clin. Epigenetics 13, 138 (2021).

19. Jia, Z. et al. Tandem CTCF sites function as insulators to balance spatial chromatin contacts and topological enhancer-promoter selection. Genome Biol. 21, 75 (2020).

20. Cockerill, P. N. Structure and function of active chromatin and DNase I hypersensitive sites. FEBS J. 278, 2182–2210 (2011).

21. Collins, B. E., Greer, C. B., Coleman, B. C. & Sweatt, J. D. Histone H3 lysine K4 methylation and its role in learning and memory. Epigenetics Chromatin 12, 7 (2019).

22. Sun, Z. et al. H3K36me3, message from chromatin to DNA damage repair. Cell Biosci. 10, 9 (2020).

23. Molina, O. et al. Epigenetic engineering reveals a balance between histone modifications and transcription in kinetochore maintenance. Nat. Commun. 7, 13334 (2016).

24. Wang, J., Lunyak, V. V. & Jordan, I. K. Chromatin signature discovery via histone modification profile alignments. Nucleic Acids Res. 40, 10642–10656 (2012).

25. Abuhashem, A., Garg, V. & Hadjantonakis, A.-K. RNA polymerase II pausing in development: orchestrating transcription. Open Biol. 12, 210220 (2022).

26. Hansen, A. S., Pustova, I., Cattoglio, C., Tjian, R. & Darzacq, X. CTCF and cohesin regulate chromatin loop stability with distinct dynamics. Elife 6, (2017).

27. Hou, Y. et al. Integrative characterization of G-Quadruplexes in the three-dimensional chromatin structure. Epigenetics 14, 894–911 (2019).

28. LaMere, S. A., Thompson, R. C., Komori, H. K., Mark, A. & Salomon, D. R. Promoter H3K4 methylation dynamically reinforces activation-induced pathways in human CD4 T cells. Genes Immun. 17, 283–297 (2016).

29. Shlyueva, D., Stampfel, G. & Stark, A. Transcriptional enhancers: from properties to genome-wide predictions. Nat. Rev. Genet. 15, 272–286 (2014).

30. Pennacchio, L. A., Bickmore, W., Dean, A., Nobrega, M. A. & Bejerano, G. Enhancers: five essential questions. Nat. Rev. Genet. 14, 288–295 (2013).

31. Zhang, B. et al. A dynamic H3K27ac signature identifies VEGFA-stimulated endothelial enhancers and requires EP300 activity. Genome Res. 23, 917–927 (2013).

32. Karmodiya, K., Krebs, A. R., Oulad-Abdelghani, M., Kimura, H. & Tora, L. H3K9 and H3K14 acetylation co-occur at many gene regulatory elements, while H3K14ac marks a subset of inactive inducible promoters in mouse embryonic stem cells. BMC Genomics 13, 424 (2012).

33. Barrera, L. O. et al. Genome-wide mapping and analysis of active promoters in mouse embryonic stem cells and adult organs. Genome Res. 18, 46–59 (2008).

34. Boix, C. A., James, B. T., Park, Y. P., Meuleman, W. & Kellis, M. Regulatory genomic circuitry of human disease loci by integrative epigenomics. Nature 590, 300–307 (2021).

35. Putiri, E. L. & Robertson, K. D. Epigenetic mechanisms and genome stability. Clin. Epigenetics 2, 299–314 (2011).

36. Sarkies, P. & Sale, J. E. Cellular epigenetic stability and cancer. Trends Genet. 28, 118–127 (2012).

37. Muntean, A. G. & Hess, J. L. Epigenetic dysregulation in cancer. Am. J. Pathol. 175, 1353–1361 (2009).

38. Dao, L. T. M. & Spicuglia, S. Transcriptional regulation by promoters with enhancer function. Transcription 9, 307–314 (2018).

39. Dao, L. T. M. et al. Genome-wide characterization of mammalian promoters with distal enhancer functions. Nat. Genet. 49, 1073–1081 (2017).

40. Andersson, R. & Sandelin, A. Determinants of enhancer and promoter activities of regulatory elements. Nat. Rev. Genet. 21, 71–87 (2020).

41. Ansari, M. Bone tissue regeneration: biology, strategies and interface studies. Prog Biomater 8, 223–237 (2019).

42. Duah, M. et al. Thymus Degeneration and Regeneration. Front. Immunol. 12, 706244 (2021).

43. Liao, Y., Wang, J., Jaehnig, E. J., Shi, Z. & Zhang, B. WebGestalt 2019: gene set analysis toolkit with revamped UIs and APIs. Nucleic Acids Res. 47, W199–W205 (2019).

44. Kanehisa, M., Sato, Y., Kawashima, M., Furumichi, M. & Tanabe, M. KEGG as a reference resource for gene and protein annotation. Nucleic Acids Res. 44, D457–62 (2016).

45. Palmer, D., Fabris, F., Doherty, A., Freitas, A. A. & de Magalhães, J. P. Ageing transcriptome meta-analysis reveals similarities and differences between key mammalian tissues. Aging 13, 3313–3341 (2021).

46. GTEx Consortium. The GTEx Consortium atlas of genetic regulatory effects across human tissues. Science 369, 1318–1330 (2020).

47. Yanai, I. et al. Genome-wide midrange transcription profiles reveal expression level relationships in human tissue specification. Bioinformatics 21, 650–659 (2005).

48. Mas, G. et al. Promoter bivalency favors an open chromatin architecture in embryonic stem cells. Nat. Genet. 50, 1452–1462 (2018).

49. Bernstein, B. E. et al. A bivalent chromatin structure marks key developmental genes in embryonic stem cells. Cell 125, 315–326 (2006).

50. Shoaib, M. et al. Histone H4 lysine 20 mono-methylation directly facilitates chromatin openness and promotes transcription of housekeeping genes. Nat. Commun. 12, 4800 (2021).

51. Buniello, A. et al. The NHGRI-EBI GWAS Catalog of published genome-wide association studies, targeted arrays and summary statistics 2019. Nucleic Acids Res. 47, D1005–D1012 (2019).

52. Simón-Sánchez, J. et al. Genome-wide association study reveals genetic risk underlying Parkinson’s disease. Nat. Genet. 41, 1308–1312 (2009).

53. Chang, D. et al. A meta-analysis of genome-wide association studies identifies 17 new Parkinson’s disease risk loci. Nat. Genet. 49, 1511–1516 (2017).

54. Streeter, I. et al. The human-induced pluripotent stem cell initiative-data resources for cellular genetics. Nucleic Acids Res. 45, D691–D697 (2017).

55. Wang, X.-L. & Li, L. Cell type-specific potential pathogenic genes and functional pathways in Alzheimer’s Disease. BMC Neurol. 21, 381 (2021).

56. Arena, G., Modjtahedi, N. & Kruger, R. Exploring the contribution of the mitochondrial disulfide relay system to Parkinson’s disease: the PINK1/CHCHD4 interplay. Neural Regeneration Res. 16, 2222–2224 (2021).

57. Kennedy, R. B. et al. Genome-wide analysis of polymorphisms associated with cytokine responses in smallpox vaccine recipients. Hum. Genet. 131, 1403–1421 (2012).

58. Suzuki, G. et al. Over-expression of a human chromosome 22q11.2 segment including TXNRD2, COMT and ARVCF developmentally affects incentive learning and working memory in mice. Hum. Mol. Genet. 18, 3914–3925 (2009).

59. Stechmiller, J. K. et al. Biobehavioral Mechanisms Associated With Nonhealing Wounds and Psychoneurologic Symptoms (Pain, Cognitive Dysfunction, Fatigue, Depression, and Anxiety) in Older Individuals With Chronic Venous Leg Ulcers. Biol. Res. Nurs. 21, 407–419 (2019).

60. Fu, Y. et al. Systematically Analyzing the Pathogenic Variations for Acute Intermittent Porphyria. Front. Pharmacol. 10, 1018 (2019).

61. Srour, M. et al. Dysfunction of the Cerebral Glucose Transporter SLC45A1 in Individuals with Intellectual Disability and Epilepsy. Am. J. Hum. Genet. 100, 824–830 (2017).

62. 1000 Genomes Project Consortium et al. A global reference for human genetic variation. Nature 526, 68–74 (2015).

63. Wang, J. et al. CAUSALdb: a database for disease/trait causal variants identified using summary statistics of genome-wide association studies. Nucleic Acids Res. 48, D807–D816 (2020).

64. Pan UKBB. https://pan.ukbb.broadinstitute.org.

65. Azevedo, T., Dimitri, G. M., Lió, P. & Gamazon, E. R. Multilayer modelling of the human transcriptome and biological mechanisms of complex diseases and traits. npj Systems Biology and Applications 7, 1–13 (2021).

66. Kupper, N., Ge, D., Treiber, F. A. & Snieder, H. Emergence of novel genetic effects on blood pressure and hemodynamics in adolescence: the Georgia Cardiovascular Twin Study. Hypertension 47, 948–954 (2006).

67. Shih, P.-A. B. & O’Connor, D. T. Hereditary determinants of human hypertension: strategies in the setting of genetic complexity. Hypertension 51, 1456–1464 (2008).

68. Davis, L. K. et al. Partitioning the heritability of Tourette syndrome and obsessive compulsive disorder reveals differences in genetic architecture. PLoS Genet. 9, e1003864 (2013).

69. Gusev, A. et al. Partitioning heritability of regulatory and cell-type-specific variants across 11 common diseases. Am. J. Hum. Genet. 95, 535–552 (2014).

70. Torres, J. M. et al. Cross-tissue and tissue-specific eQTLs: partitioning the heritability of a complex trait. Am. J. Hum. Genet. 95, 521–534 (2014).

71. Bulik-Sullivan, B. K. et al. LD Score regression distinguishes confounding from polygenicity in genome-wide association studies. Nat. Genet. 47, 291–295 (2015).

72. Finucane, H. K. et al. Partitioning heritability by functional annotation using genome-wide association summary statistics. Nat. Genet. 47, 1228–1235 (2015).

73. Gazal, S. et al. Combining SNP-to-gene linking strategies to identify disease genes and assess disease omnigenicity. Nat. Genet. (2022) doi:10.1038/s41588-022-01087-y.

74. Shi, C. et al. A quantitative discriminant method of elbow point for the optimal number of clusters in clustering algorithm. Eurasip J. Wirel. Commun. Network. 2021, 1–16 (2021).

75. Chatterjee, S. & Ahituv, N. Gene Regulatory Elements, Major Drivers of Human Disease. Annu. Rev. Genomics Hum. Genet. 18, 45–63 (2017).

76. Matharu, N. K. & Ahanger, S. H. Chromatin Insulators and Topological Domains: Adding New Dimensions to 3D Genome Architecture. Genes 6, 790–811 (2015).

77. McArthur, E. & Capra, J. A. Topologically associating domain boundaries that are stable across diverse cell types are evolutionarily constrained and enriched for heritability. Am. J. Hum. Genet. 108, 269–283 (2021).

78. Xuan, Z., Zhao, F., Wang, J., Chen, G. & Zhang, M. Q. Genome-wide promoter extraction and analysis in human, mouse, and rat. Genome Biol. 6, R72 (2005).

79. Panigrahi, A. & O’Malley, B. W. Mechanisms of enhancer action: the known and the unknown. Genome Biol. 22, 108 (2021).

80. Brown, T. A. The Human Genome. (Wiley-Liss, 2002).

81. Pedregosa, F. et al. Scikit-learn: Machine Learning in Python. arXiv [cs.LG] (2012).

82. Feng, J., Liu, T., Qin, B., Zhang, Y. & Liu, X. S. Identifying ChIP-seq enrichment using MACS. Nat. Protoc. 7, 1728–1740 (2012).

83. Hinrichs, A. S. et al. The UCSC Genome Browser Database: update 2006. Nucleic Acids Res. 34, D590–8 (2006).

84. Abugessaisa, I. et al. refTSS: A Reference Data Set for Human and Mouse Transcription Start Sites. J. Mol. Biol. 431, 2407–2422 (2019).

85. Quinlan, A. R. & Hall, I. M. BEDTools: a flexible suite of utilities for comparing genomic features. Bioinformatics 26, 841–842 (2010).

86. Howe, K. L. et al. Ensembl 2021. Nucleic Acids Res. 49, D884–D891 (2021).

87. Abugessaisa, I. et al. FANTOM5 CAGE profiles of human and mouse reprocessed for GRCh38 and GRCm38 genome assemblies. Sci Data 4, 170107 (2017).

88. Roadmap Epigenomics Consortium et al. Integrative analysis of 111 reference human epigenomes. Nature 518, 317–330 (2015).

89. Zhang, Z. et al. HeRA: an atlas of enhancer RNAs across human tissues. Nucleic Acids Res. 49, D932–D938 (2021).

90. Duttke, S. H., Chang, M. W., Heinz, S. & Benner, C. Identification and dynamic quantification of regulatory elements using total RNA. Genome Res. 29, 1836–1846 (2019).

91. Harrow, J. et al. GENCODE: the reference human genome annotation for The ENCODE Project. Genome Res. 22, 1760–1774 (2012).

92. Pruim, R. J. et al. LocusZoom: regional visualization of genome-wide association scan results. Bioinformatics 26, 2336–2337 (2010).

93. Pohl, A. & Beato, M. bwtool: a tool for bigWig files. Bioinformatics 30, 1618–1619 (2014).

94. Köhler, S. et al. The Human Phenotype Ontology in 2021. Nucleic Acids Res. 49, D1207–D1217 (2021).

95. Zhao, J. H. gap: Genetic Analysis Package. J. Stat. Softw. 23, 1–18 (2008).

