## Supplementary Figures for "Minimum entropy framework identifies a novel class of genomic functional elements and reveals regulatory mechanisms at human disease loci"

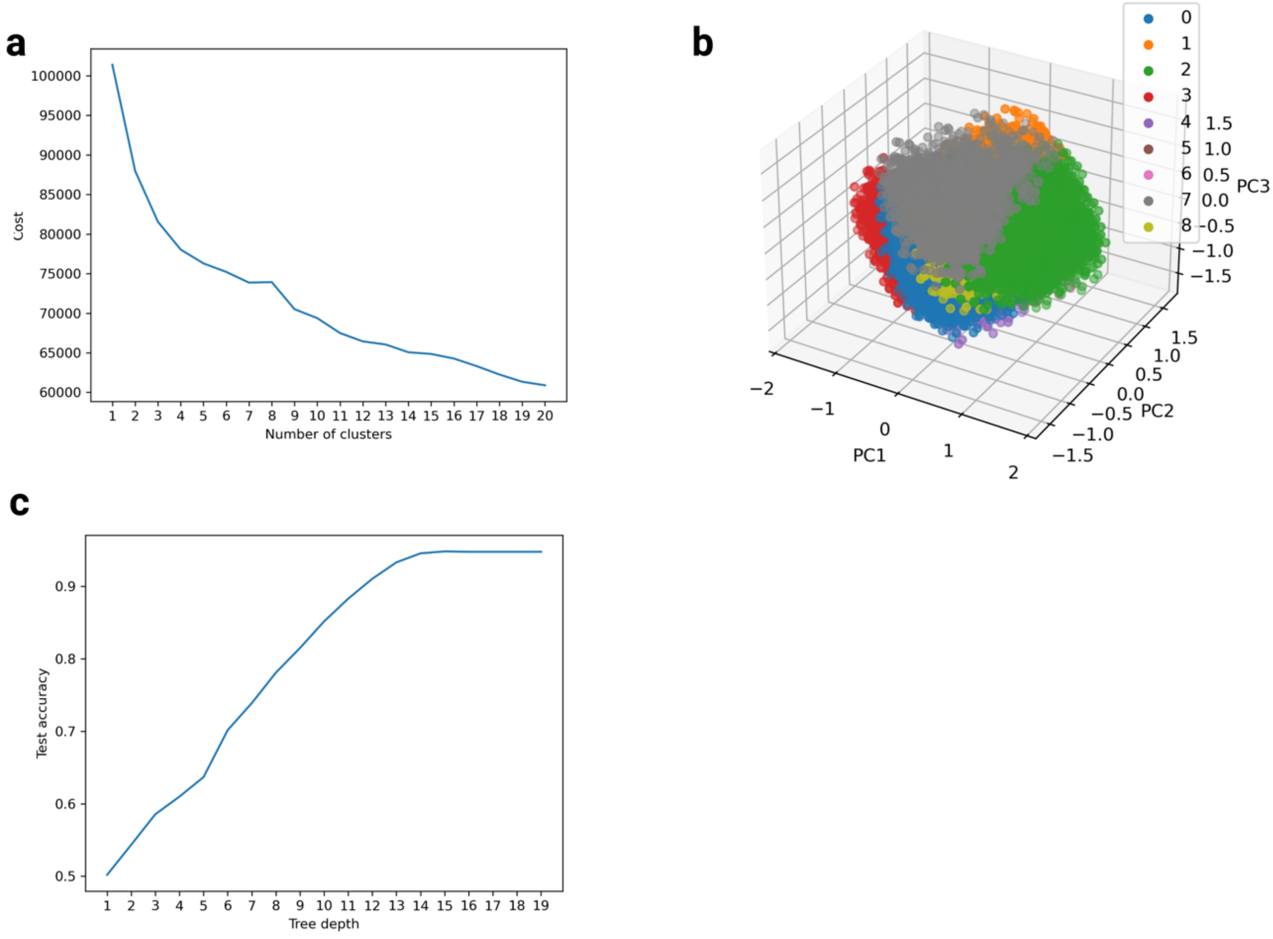


***Supplementary Figure 1. CoRE-BED decision tree training. a,*** *K-modes clustering cost plotted as a function of the number of clusters (1-19) evaluated. We selected 9 as the ideal number of clusters representing the data, as the cost was not significantly reduced beyond this point.* ***b,*** *Each of the unique putative regulatory elements from the decision tree training and test sets was plotted using the first three principal components. Labels were derived by performing k-modes clustering using the previously-learned* $k = 9$*.* ***c,*** *CoRE-BED decision tree accuracy in the test set as a function of maximum tree depth. The best-performing decision tree used a maximum depth of 15.*

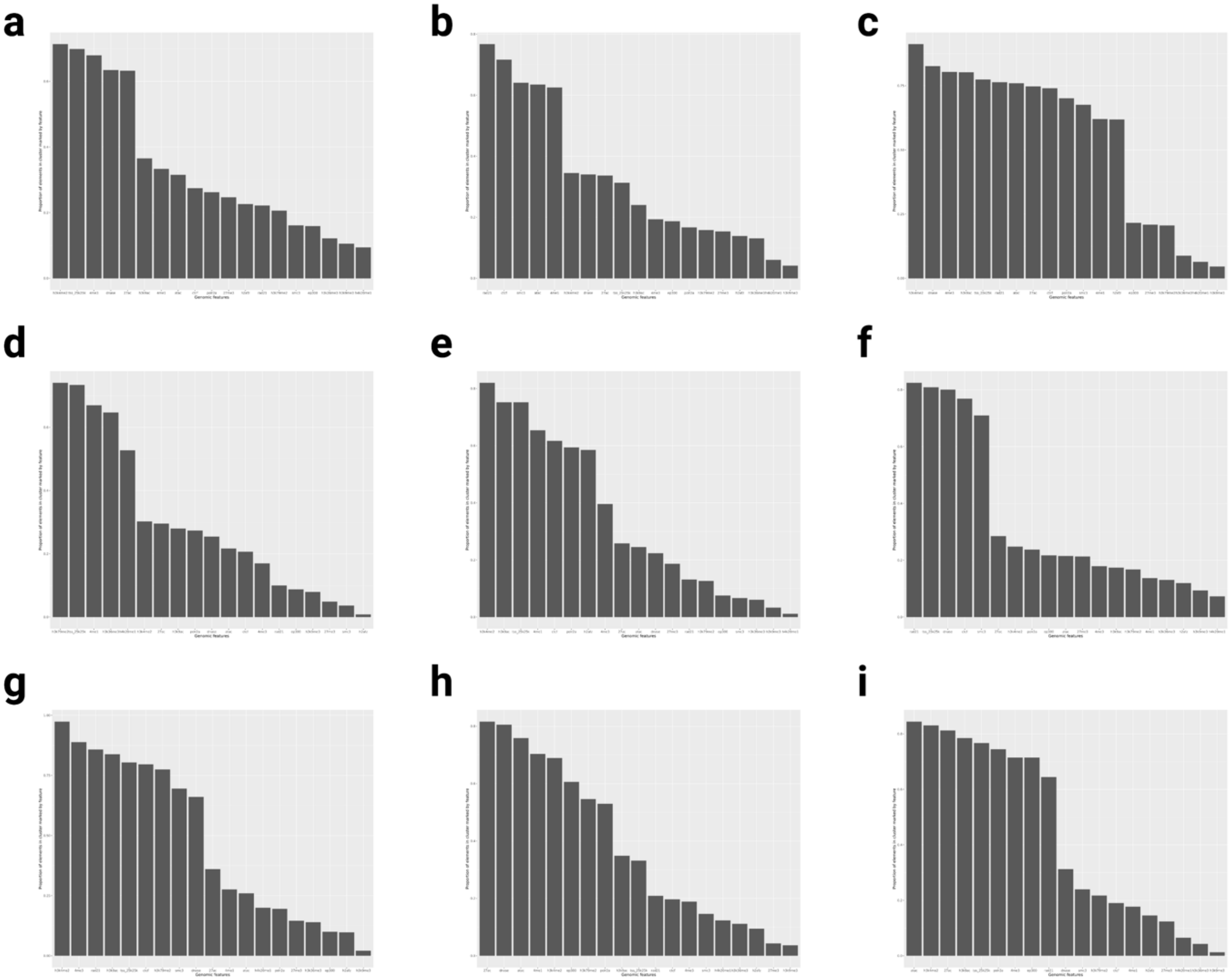


***Supplementary Figure 2. Enrichment of genomic features in each of the 9 learned CoRE-BED functional categories. a,*** *Class 0 (Non-functional).* ***b,*** *Class 1 (distal insulator).* ***c,*** *Class 2 (active promoter 1).* ***d,*** *Class 3 (transcribed gene body).* ***e,*** *Class 4 (DAE).* ***f,*** *Class 5 (proximal insulator).* ***g,*** *Class 6 (active promoter 2).* ***h,*** *Class 7 (active enhancer).* ***i,*** *Class 8 (ePromoter).*

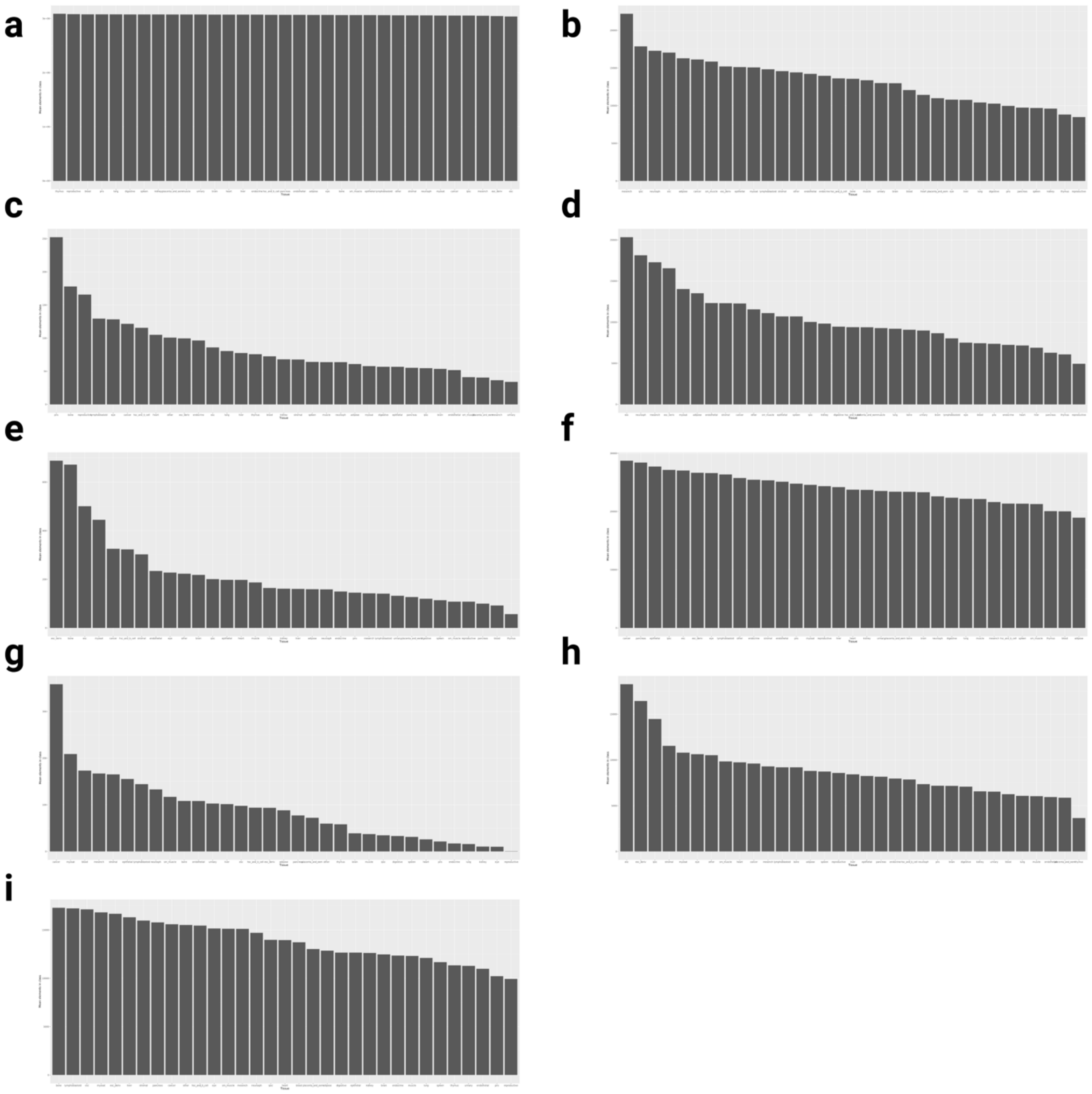

***Supplementary Figure 3. By-tissue enrichments of the 9 CoRE-BED functional classes, ordered from highest to lowest. a,*** *Class 0 (Non-functional).* ***b,*** *Class 1 (distal insulator).* ***c,*** *Class 2 (active promoter 1).* ***d,*** *Class 3 (transcribed gene body).* ***e,*** *Class 4 (DAE).* ***f,*** *Class 5 (proximal insulator).* ***g,*** *Class 6 (active promoter 2).* ***h,*** *Class 7 (active enhancer).* ***i,*** *Class 8 (ePromoter).*

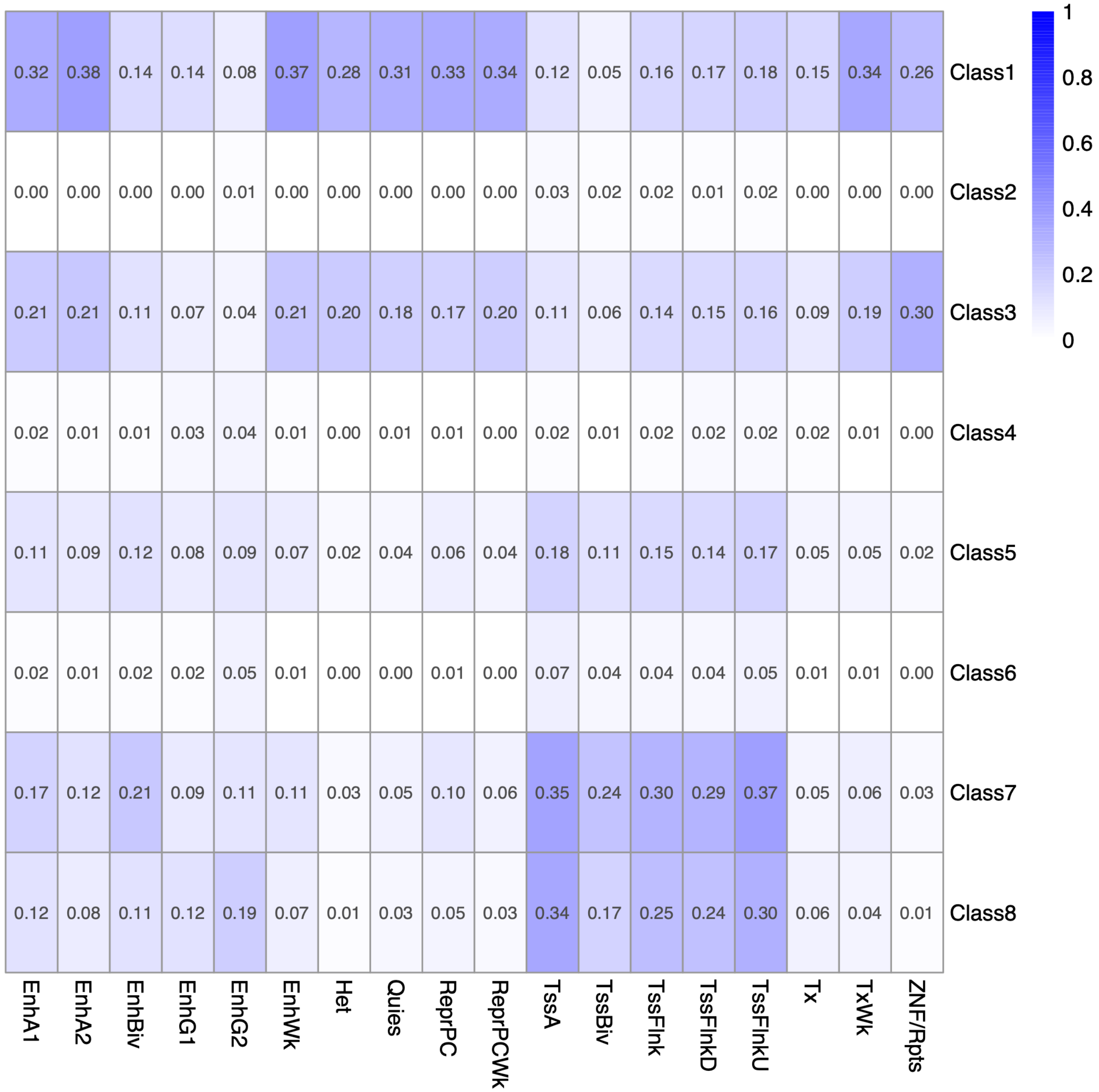


***Supplementary Figure 4. Pairwise comparison of CoRE-BED functional classes with ChromHMM chromatin states.*** *Pairwise Jaccard similarity indices were computed to determine the degree of similarity of elements in each CoRE-BED functional class to those in each ChromHMM chromatin state.*

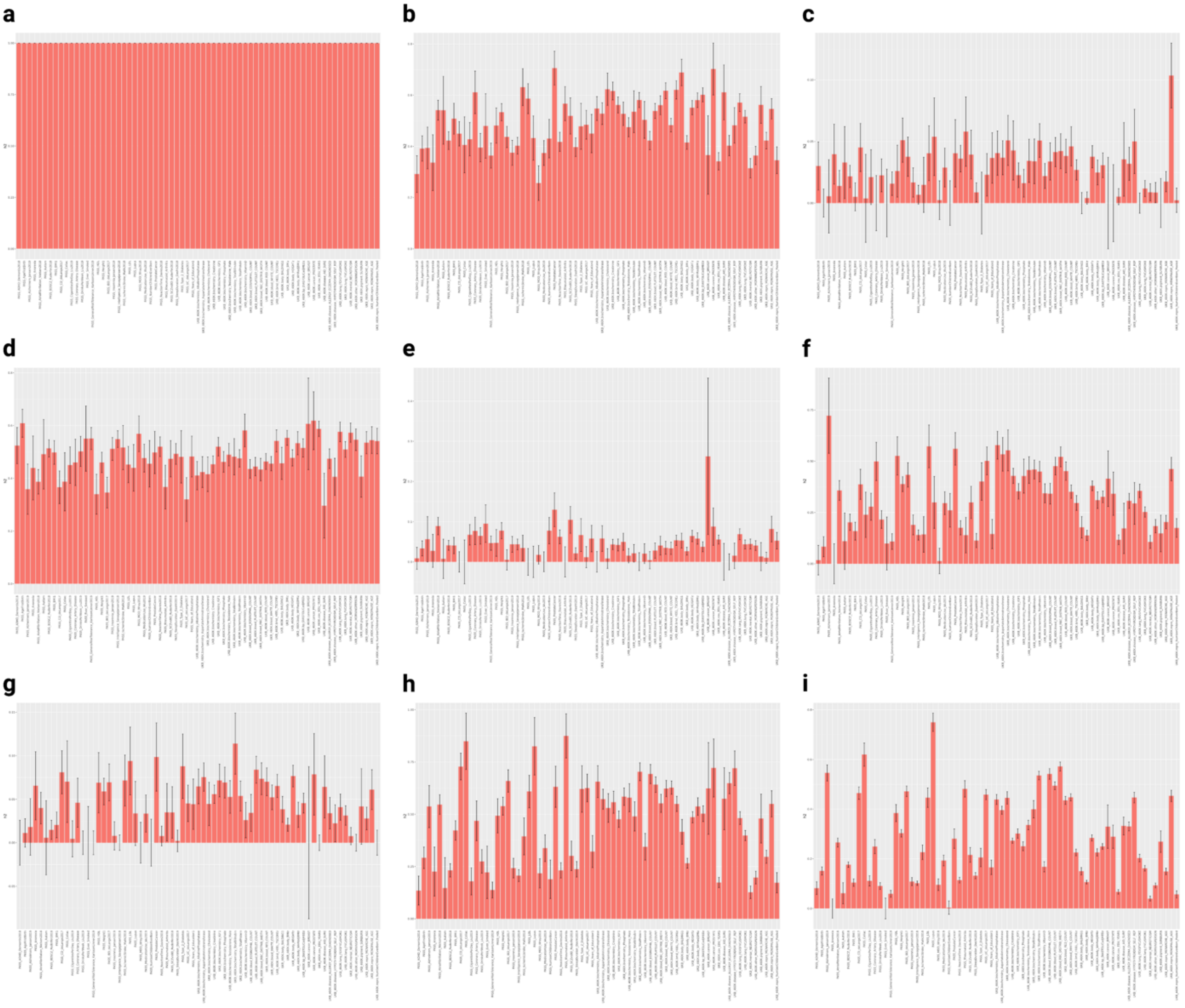


***Supplementary Figure 5. Estimated proportions of SNP heritability across 69 GWAS that are captured by CoRE-BED functional predictions. a,*** *Any CoRE-BED functional prediction.* ***b,*** *Class 1 (distal insulator).* ***c,*** *Class 2 (active promoter 1).* ***d,*** *Class 3 (transcribed gene body).* ***e,*** *Class 4 (DAE).* ***f,*** *Class 5 (proximal insulator).* ***g,*** *Class 6 (active promoter 2).* ***h,*** *Class 7 (active enhancer).* ***i,*** *Class 8 (ePromoter). Error bars represent standard error.*
