## Supplementary Table Captions for "Minimum entropy framework identifies a novel class of genomic functional elements and reveals regulatory mechanisms at human disease loci"

**Supplementary Table 1. Transcription factor binding motifs enriched in CoRE-BED ePromoters.** We started with a list of genomic regions predicted by CoRE-BED to be a Class 8 element (ePromoter) in at least one cell or tissue type. Within these regions, we found 130 transcription factor binding motifs that were only significantly (Benjamini-Hochberg-corrected $p < 0.05$) enriched in ePromoters and not within the remaining genome.

**Supplementary Table 2. DAEs per tissue and DAE-regulated genes per tissue.** The first tab contains mean values of Class 4 elements (DAEs) across all 33 CoRE-BED cell and tissue types. The second tab contains total counts of the number of unique genes predicted to be under the control of a burst element in each tissue type.

**Supplementary Table 3. GWAS Catalog SNP associations in a burst element.** This table contains the 1,697 GWAS Catalog SNPs that fall within a CoRE-BED burst element in at least one cell or tissue type. The columns on the far right denote in which of the 33 major cell and tissue types each was predicted to be a burst element.

**Supplementary Table 4. Astrocyte eQTLs within astrocyte-specific CoRE-BED burst elements.** The first tab contains all 126 mapped astrocyte eQTLs that fall with an astrocyte-specific burst element. The second tab contains a list of the 86 unique genes linked to these eQTLs.

**Supplementary Table 5. Partitioned heritability estimates for 69 GWAS included in the cS2G paper.** The first tab contains the proportions of SNP heritability from each GWAS captured by CoRE-BED functional classes. The second tab contains absolute values of SNP heritability for each trait captured by each CoRE-BED class.
